# Aβ-affine bifunctional copper chelators capable of Aβ-induced oxidative stress reduction and amyloid disaggregation

**DOI:** 10.1101/2023.10.10.561649

**Authors:** Olga Krasnovskaya, Daniil Abramchuk, Alexander Vaneev, Peter Gorelkin, Maxim Abakumov, Roman Timoshenko, Nelly Chmelyuk, Veronika Vadekhina, Regina Kuanaeva, Evgeniy V. Dubrovin, Vasilii Kolmogorov, Elena Beloglazkina, Alexander Erofeev

## Abstract

Alz-5 acts as a bifunctional chelator that can interact with various Aβ aggregates and reduce their neurotoxicity. Single-cell ROS measurements provided by Pt-nanoelectrode technique revealed a significant antioxidant properties of Alz-5 in Aβ_42_ - affected SH-SY5Y cell. AFM data obtained on Aβ_42_ fibrils clearly indicate an anti-aggregating property of Alz-5. Young’s modulus mapping on living SH-SY5Y cells revealed an ability of Alz-5 to decrease cell rigidity in Aβ_42_ - affected SH-SY5Y cells.

---

Alzheimer’s disease (AD) is the most common neurodegenerative disorder, which is characterized by intracerebral β-amyloid (Aβ) aggregation, τ-hyperphosphorylation, and loss of cholinergic neurons. The other important hallmarks of AD are oxidative stress, metal dyshomeostasis and inflammation.^1^

A breakdown in metal homeostasis is an important therapeutic target used in the development of anti-AD drugs.^2^ AD-affected brains are associated with high concentrations of copper, iron, and zinc in amyloid. Metal cations can stabilize neurotoxic soluble Aβ oligomers and promote reactive oxygen species (ROS) formation.^3^ Also, soluble Aβ oligomers instead of insoluble Aβ plaques were found to be the most neurotoxic species, capable of synapse loss and neuronal injury affection.^6,7^ Oxidative stress is one important consequence of metal ion dyshomeostasis. In particular, the redox-activity of the Cu(I/II) pair results in ROS overproduction from Fenton-like reactions.^8,9^ Given the multifactorial etymology of AD, the design of several-targeted drugs is of high interest. Thus, imparting antioxidant or metal-binding functions to drugs that are clinically available is also on demand.^10,11^

The use of metal chelation agents is a promising AD-treatment strategy. Metal chelators can reduce Aβ aggregation, ROS formation, and neurotoxicity in vitro.^12^ Bifunctional chelators (BFCs), consisting of an Aβ-affine aromatic moiety, and a metal chelating scaffold are widely used frameworks for the design of diagnostic agents for positron emission tomography (PET) Aβ plaques visualization. ^13-19^

Also, several studies have proven the therapeutic potential of such BFCs, via demonstrating their ability to modulate amyloid aggregation, reduce amyloid toxicity and oxidative stress.^20-23^ Recently, we summarised the use of BFCs and coordination compounds based on them for therapy and imaging of AD.^24^

Herein, we report a synthesis investigation of five bifunctional copper chelators, capable of metal-Aβ interaction disruption, based on benzothiazole and stilbene moieties. Those well-known scaffolds originated from the Pittsburgh compound B (PiB), a radioactive analogue of Aβ staining agent Thioflavin T(ThT), which is used for Aβ plaque imaging, and ^18^F-labeled stilbene derivative Florbetaben, which is also a PET radiotracer for Aβ-amyloid imaging.^25,26^.

As copper-chelating scaffold, thiocarbahydrazone and thiosemicarbazone moieties were used, which were also successfully utilized in the design of amyloid-imaging drugs, as well as Cu-ATSM drugs.^27^ The choice of allyl, benzyl, and phenylethyl moieties in molecular design was guided by our early studies of the antitumor activity of copper-containing coordination compounds.^28,29^

BFCs based on benzothiazole scaffold with thiocarbahydrazone chelating moieties **Alz-1 -Alz-3** were synthesised in six steps (Scheme S1,†), BFCs based on stilbene scaffold with thiosemicarbazone chelating moieties **Alz-4, Alz-5** were synthesised in nine successful steps (Scheme S2,†). The ligands were characterized by NMR, high resolution mass spectrometry, and IR, and the purity of **Alz-1-Alz-5** was confirmed by HPLC (Figures S1-S37,†).

Since BFCs are intended to act as copper chelating agents, their cell toxicity in the presence of copper cations should not be significant. Thus, the toxicity of the drugs on cancer derived liver (HepG2) and brain (SH-SY5Y) cells was assessed. Based on the results (Figure S38,S39, †), **Alz-5** was the least toxic compound.

Since Cu^2+^−Aβ species proved to be neurotoxic,^30,31^ the influence of metal-chelating BFCs **Alz 1-5** on Aβ_40_-induced toxicity on viability of neuroblastoma cells in the presence of monomeric Aβ_40_, as well as in the presence of preformed Aβ_40_ fibrils was accessed on human neuroblastoma cells (SH-SY5Y). The effect of copper cations on neurotoxicity, which is of obvious interest, has also been assessed. The results indicate the undoubted binding of **Alz 1-5** drugs with copper cations, as well as their ability to modulate Aβ aggregation (Figure 2). An increased neurotoxicity of Aβ_40_ species in the presence of Cu^2+^ and **Alz 1-5** is provided by the formation of soluble Aβ_40_ oligomers of various sizes. This result is in good accordance with those obtained by Sharma et al., an increased cellular toxicity of Aβ_42_ oligomers in the presence of both Cu^2+^ and BFCs was observed._21_ **Alz 1-5** drugs showed the ability to modulate the aggregation of both amyloids and fibrils in the presence of copper cations, while not having a toxic effect in the absence of copper cations.

**Figure 1.**
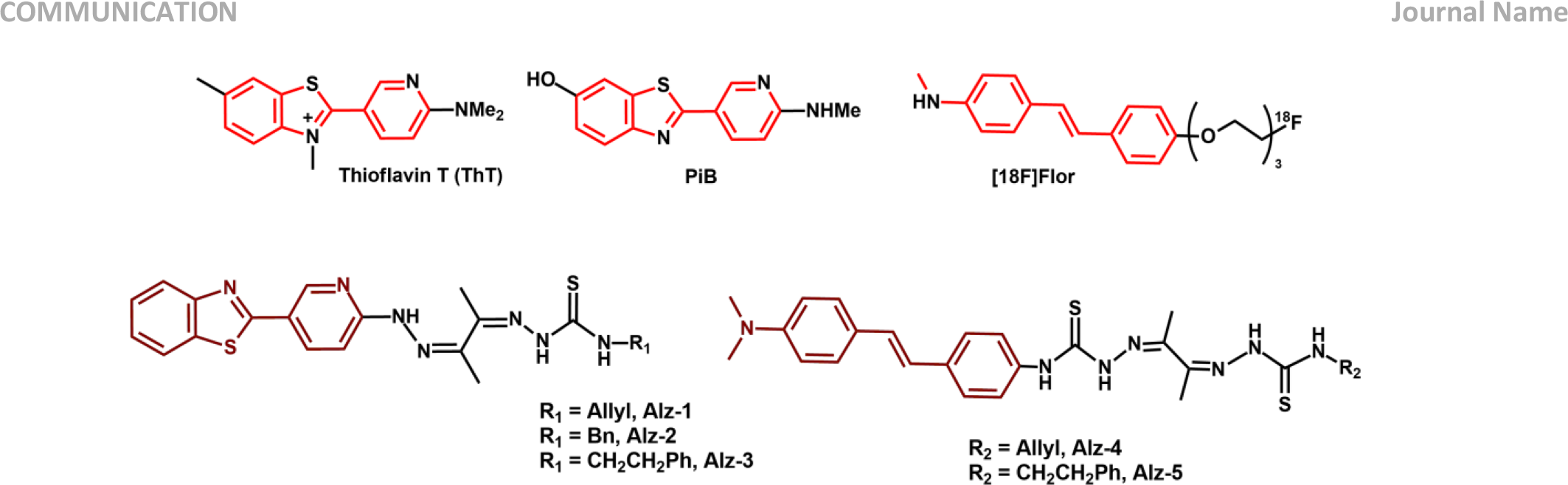
Structures of Thioflavin-T, Pittsburg compound B, ^18^– F –Florbetaben, and bifunctional compounds Alz-1 -Alz-5.

**Figure 2.**
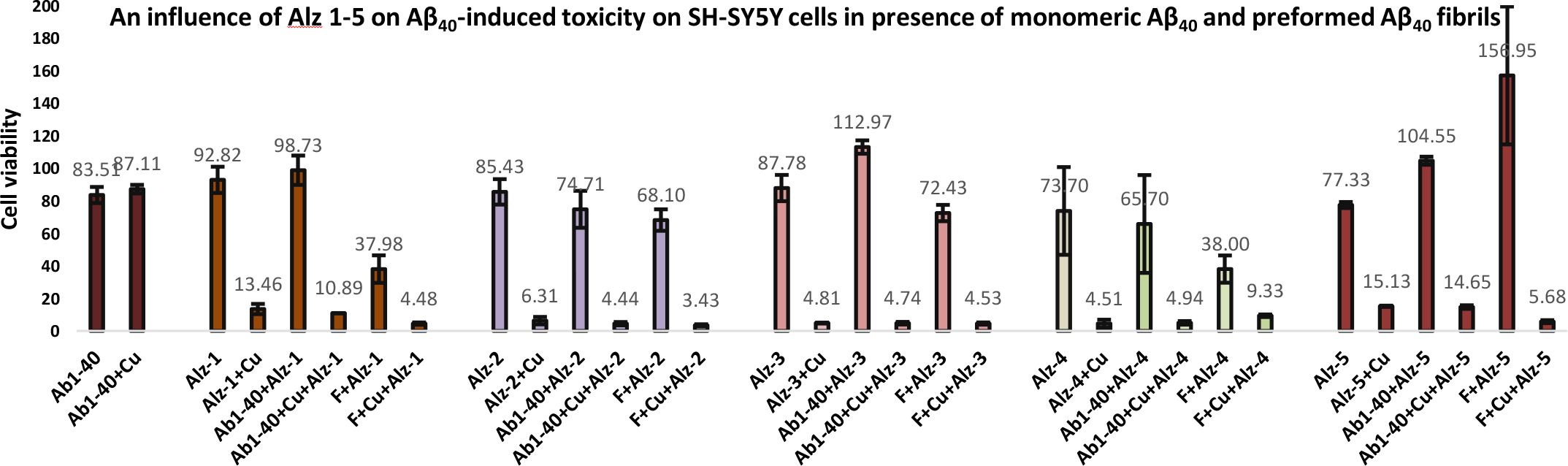
An influence of Alz 1-5 on Aβ40-induced neurotoxicity on SH-SY5Y cells in presence of monomeric Aβ40 and preformed Aβ40 fibrils. Drug (20 mM), Drug (20 mM) + CuCl_2_, Aβ40 + Drug (20 mM), Aβ40 + Drug (20 mM) + CuCl_2_, Aβ40 fibrils + Drug (20 mM), Aβ40 fibrils + Drug (20 mM) + CuCl_2_.

It is well known that the formation of Aβ aggregates is accompanied by a disruption in mitochondrial activity and changes in the level of ROS, which causes multifaced toxicity.^32^ Thus, drugs that specifically scavenge oxygen radicals may have a particular therapeutic efficacy.^33^ The ability of **Alz-5** to reduce oxidative stress caused by the presence of Aβ amyloids was of extremely interest. To assess intracellular ROS level, single-cell amperometric ROS measurement using Pt-nanoelectrodes was used.^34,35^ The effect of **Alz-5** on the intracellular ROS levels in the presence of Aβ_42_ in single SH-SY5Y cells was accessed (Figure 3).

**Figure 3.**
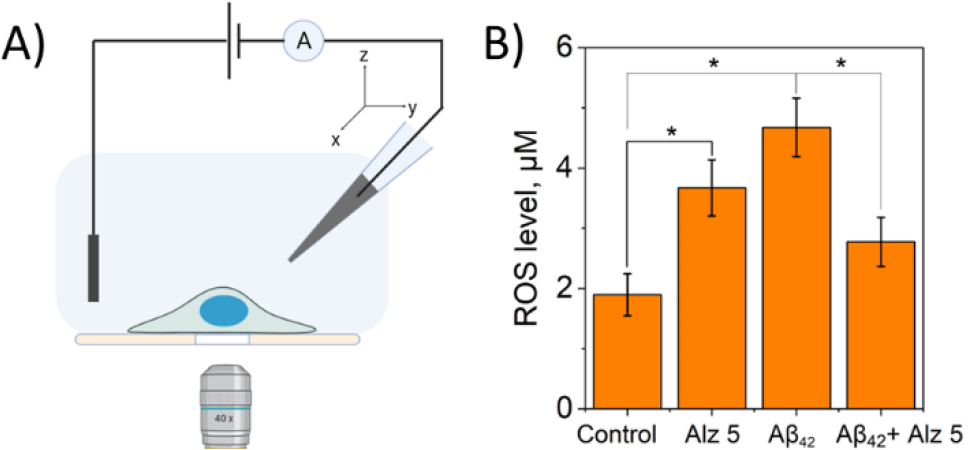
Electrochemical intracellular ROS measurement (A) Scheme of measurement (B) The ROS level inside SH-SY5Y cells after simultaneous incubation with Aβ_42_ and Alz 5. The results are shown as mean, SE and (*) p < 0.05 (one-way ANOVA)

The intracellular level of ROS increased by 2.2 times after incubation with Aβ_42_. Even though **Alz-5** caused an increase in intracellular ROS level, simultaneous incubation of Aβ_42_ with **Alz-5** leads to a decrease in intracellular ROS level compared to Aβ_42_ 1.7-fold, thereby confirming the antioxidant activity of **Alz-5** and its ability to reduce Aβ-induced oxidative stress. Previously, Sun et al. reported an amphiphilic BFC with antioxidant activity, which was confirmed by ascorbate consumption assays.^22^ Herein is the first example of real-time in-cell monitoring of Aβ-induced oxidative stress reduction, caused by bifunctional copper chelator.

To confirm the ability of drugs to modulate amyloid disaggregation, the effects of **Alz-5** drug on amyloids which can be observed with the «naked eye» were of great interest. The direct effect of **Alz-5** drug on the mechanistic properties of amyloids would be irrefutable evidence of its disaggregation ability. To date, several TEM images of the inhibition of Aβ_42_ aggregation by BFCs have been provided. Herein, to evaluate amyloid disaggregation process atomic force microscopy (AFM) was used due to its high resolution.

The aggregation of Aβ_42_ amyloid for 24 hours at 37°C leads to well-defined Aβ_42_ fibrils, aggregates ∼300 nm length, as confirmed by AFM images (Figure 4, A). A metal-mediated aggregation was observed after CuCl_2_ addition, along with the extension of aggregates up to 400 nm (Figure 4, B). The addition of **Alz-5** drug leads to a shortening of aggregates up to ∼150 nm, along with a decrease in tendency to aggregate (Figure 4, C). At the same time, in the presence of both copper chloride and **Alz-5**, no visible fibrils were detected. Since in the presence of both copper chloride and **Alz-5** no fibrils were observed, hoverer, the diameter of the aggregates was calculated. Summarized data on fibril length estimated using AFM are presented in Figure 4, E.

**Figure 4.**
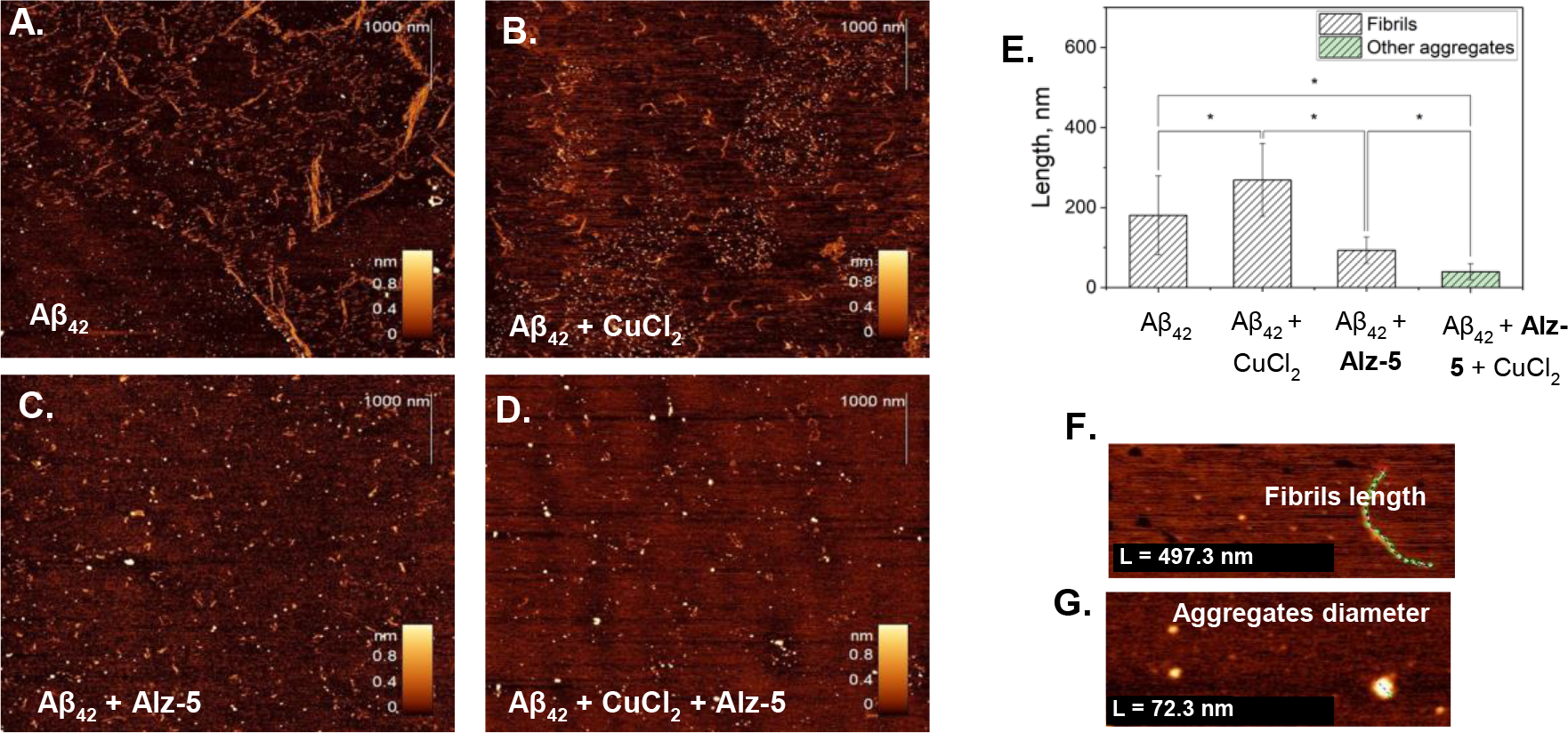
Diverse Ab_42_ assemblies imaged by AFM with data analysis. AFM images of Aβ_42_ fibrils (A); Aβ_42_ + CuCl_2_ (B); Aβ_42_ + Alz-5(C); Aβ_42_ + CuCl_2_ + Alz-5(D); E) Histogram of Aβ_42_ aggregates’ size (fibrils and oligomers). The error bars represent standard deviation (N_Aβ42_ = 214, N_Aβ42+CuCl2_ = 135, N_Aβ42+Alz-5_ = 157, N_Aβ42+CuCl2+Alz-5_ = 62). The asterisk (*) indicate significant differences, p < 0.05, one-way ANOVA; Example of measured fibril length (F) and aggregates diameter (G).

Thus, AFM data clearly indicate the anti-aggregating properties of **Alz-5** both in the absence and presence of Cu^2+^. Amyloid fibrils have been repeatedly reported to cause cytoskeleton modifications, such as microtubule disassembly or actin polymerization, which results in degeneration in neurons.^36-38^ The presence of amyloids in neuronal cells can lead to structural reorganization, a significant increase in fibrillar actin leads to increased cellular rigidity, which can be estimated with scanning ion-conductance microscopy (SICM) technique.^39,40^ It was of great interest to evaluate the effect of **Alz-5** on the stiffness of a neuronal cell exposed to amyloid. Thus, we provided young’s modulus mapping of Aβ_42_ aggregates formation on living SH-SY5Y cell surfaces after incubation with **Alz-5** only, Aβ_42_ and Aβ_42_ + **Alz-5** (Figure 5, A). Expectedly, the addition of Aβ_42_ to SH-SY5Y cells resulted in a dramatic increase in cell stiffness, which was inhibited by the addition of **Alz-5**. The decrease in Young’s modulus can be explained by the destruction of amyloid, which occurs even in the absence of copper cations. Summarized data on Young modulus value estimated using SICM are presented in Figure 5, B. Despite the incubation of SH-SY5Y cells with **Alz-5** only led to an increase of Young’s modulus in comparison with control cells, the obvious drop in cell rigidity caused by amyloid after **Alz-5** leaves no doubt about the effectiveness of **Alz-5** as an anti-aggregating agent that acts even in the absence of copper cations.

**Figure 5.**
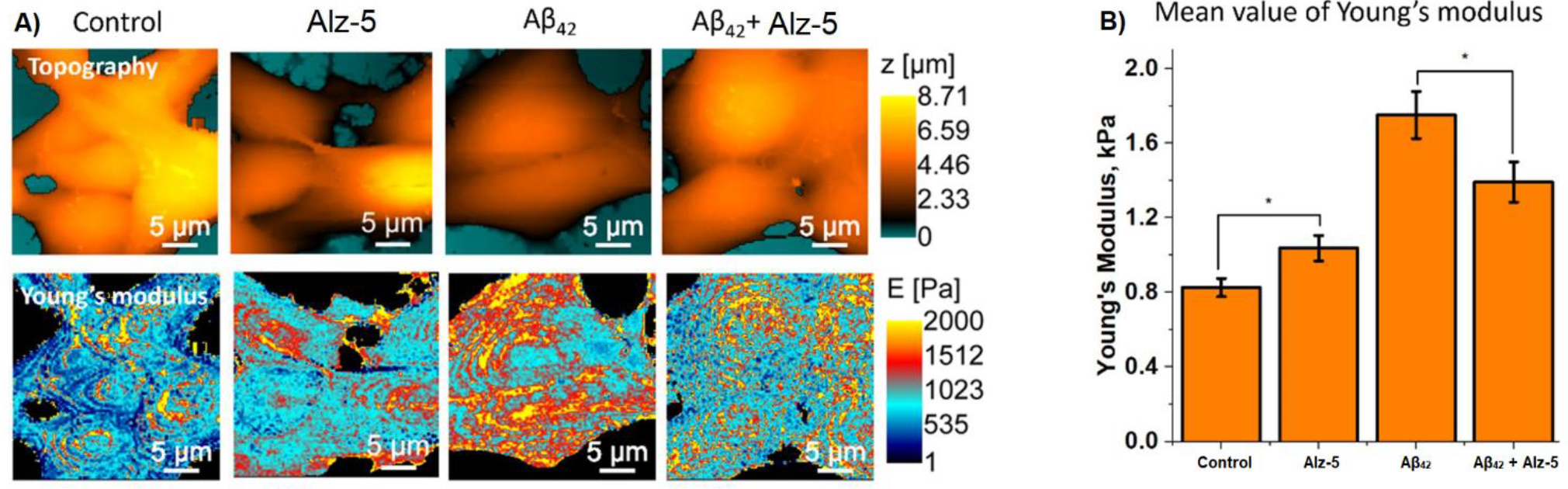
Topography and Young’s modulus maps of control SH-SY5Y cells, after incubation with Alz-5, Aβ42 and Aβ42 + Alz-5 (A); B – Mean value of Young’s modulus (*p≤0.01, one-Way ANOVA) (B).

Herein, we report design and synthesis of BFCs **Alz-1 -Alz-5**, capable of both Aβ_40-42_ binding and metal chelation. **Alz-5** showed a promising ability to modulate the aggregation of both Aβ_40_ amyloids and fibrils in the presence of copper cations. Single-cell ROS measurements provided by the Pt-nanoelectrode technique revealed a significant Aβ_42_-induced oxidative stress reduction in the presence of **Alz-5**. AFM data obtained on Aβ_42_ fibrils clearly indicate an anti-aggregating property of **Alz-5** both in the presence and absence of Cu^2+^. Young’s modulus mapping and confocal imaging of Aβ_42_ aggregate formation on living SH-SY5Y cells revealed a decrease in Aβ_42_ - affected SH-SY5Y cell rigidity.

The study was performed employing a unique scientific facility “Scanning ion-conductance microscope with a confocal module” (registration number 2512530) and was financially supported by the Ministry of Education and Science of the Russian Federation, Agreement No. 075-15-2022-264.

## Supporting information

Spectra&Methods

