## Supplementary material for "Aβ-affine bifunctional copper chelators capable of Aβ-induced oxidative stress reduction and amyloid disaggregation": Spectra&Methods

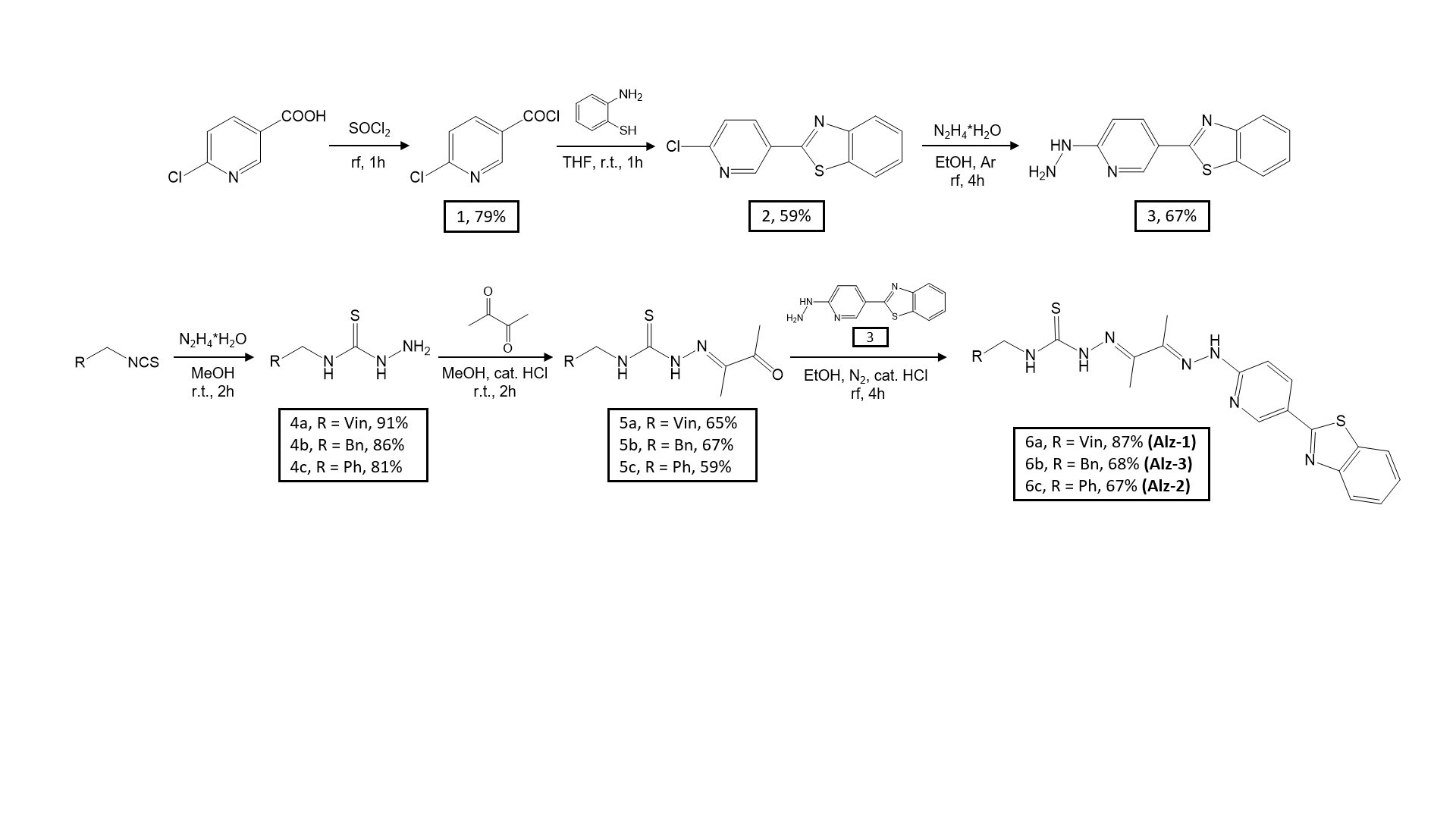

Scheme S1. Scheme of synthesis Alz-1-3

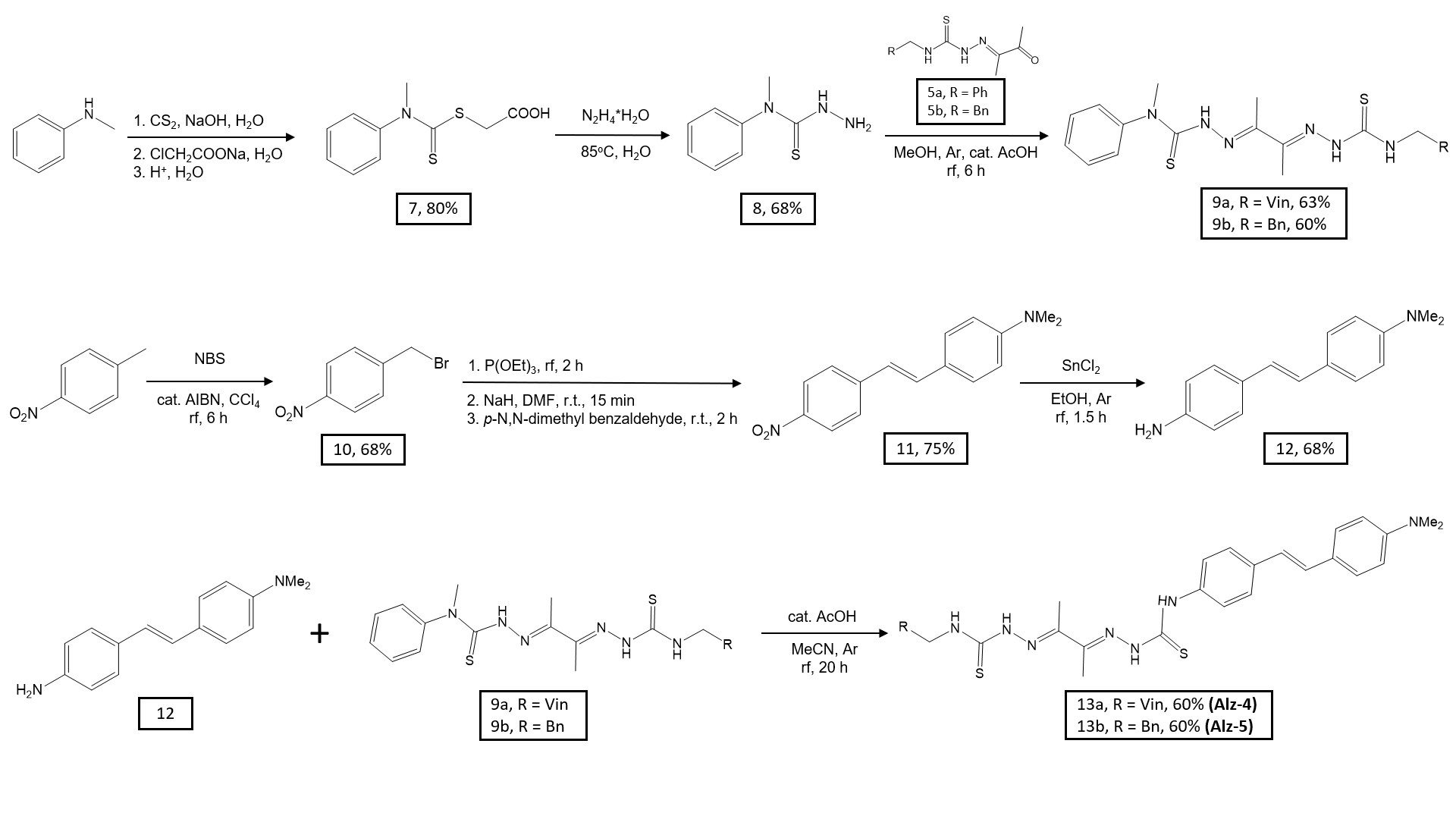

Scheme S2. Scheme of synthesis Alz-4-5

### Synthetic procedures

**Synthesis of 6-chloronicotinyl chloride (1).**

6-Chloronicotinic acid (500 mg, 3.174 mmol) was refluxed in excess thionyl chloride (5 ml) under argon for 1 hour. The mixture was then cooled to room temperature and evaporated on a rotor. 547 mg (98%) of 6-chloronicotinyl chloride was obtained in the form of a light-yellow crystalline substance.

^1^H NMR (400 MHz, CDCl_3_) δ 9,07(d, J = 1,96 Hz, 1H), 8,31 (m, 1Н), 7,51 (d, 1H, J = 7,83 Hz).

**Synthesis of 2-(6-chloropyridin-3-yl) benzothiazole (2).**

6-Chloronicotinyl chloride 1 (547 mg, 3.111 mmol) was dissolved in THF (5 ml), 2-aminothiophenol (389 mg, 3.111 mmol) was added dropwise over 10-15 minutes and then the reaction mixture was stirred at room temperature for an hour. The reaction mixture was then diluted with CH_2_Cl_2_ (5 ml) and neutralized with saturated NaHCO_3_ solution. The organic layer was separated, and the aqueous layer was washed with CH_2_Cl_2_. The organic fractions were combined, dried over Na_2_SO_4_ and evaporated on a rotor. The resulting residue was purified by flash chromatography (CH_2_Cl_2_ served as the eluent). 464 mg (59%) of 2-(6-chloropyridin-3-yl) benzothiazole 2 were obtained as a light-yellow solid.

^1^H NMR (400 MHz, CDCl_3_) δ 9.07 (d, J = 2.45 Hz, 1H), 8.36 (dd, J1 = 8.31 Hz, J2 = 2.45 Hz, 1H), 8.10 (d, J = 8.8 Hz, 1H), 7.95 (d, J = 7.82, 1H), 7.55 (m, 1H), 7.47 (m, 2H).

**Synthesis of 2-(6-hydrazinylpyridin-3-yl) benzothiazole (3).**

2-(6-Chloropyridin-3-yl) benzothiazole 2 (200 mg, 0.81 mmol) and hydrazine monohydrate (2 ml) were boiled in ethanol (10 ml) under argon for 4 hours. When cooled to room temperature, a light-yellow precipitate formed, which was filtered, washed with ethanol and ether, and left to air dry. 131 mg (67%) of 2-(6-hydrazinylpyridin-3-yl) benzothiazole 3 was obtained as a white solid (131 mg, 67%).

^1^H NMR (400 MHz, DMSO-d6) δ 8.67 (s, 1H), 8.28 (br s, 1H), 8.09- 8.05 (m, 2H), 7.94 (d, J = 7.82 Hz, 1H), 7.48 (t, J = 7.82 Hz, 1H), 7.37 (m, 1H), 6.83 (d, J = 8.8 Hz, 1H), 4.38 (br s, 2H).

**Synthesis of N-substituted hydrazinecarbothioamides (4a-c).**

General technique:

The original isothiocyanate (1 molar equivalent) was dissolved in MeOH (6.6 ml per 100 mg of solution) and stirred with hydrazine hydrate at room temperature until the isothiocyanate spot disappeared on the TLC of the reaction mixture. The solvent was then evaporated under reduced pressure. The resulting residue was washed with ether and dried in air.

**Synthesis of N-allylhydrazinecarbothioamide (4a).**

From 2.026 g (0.02 mol) of allyl isothiocyanate, as a result of the reaction with hydrazine hydrate, 2.1 g gave N-allylhydrazinecarbothioamide 4a in the form of white needle-shaped crystals. Yield: 2.43 g (91%).

^1^H NMR (400 MHz, CDCl_3_) δ 8.07 (br s, 1H), 7.52 (s, 1H), 5.81-6.00 (m, 1H), 5.09 – 5.27 (m, 2H), 4.27 (t, J = 5.7 Hz, 2H), 3.81 (br s, 2H).

^13^C NMR (101 MHz, CDCl_3_) δ 182.14, 133.84, 116.77, 46.41.

**Synthesis of N-phenylethylhydrazinecarbothioamide (4b).**

From 1 g (0.0056 mol) of 2-phenylethyl isothiocyanate, as a result of the reaction with hydrazine hydrate, 0.575 g yielded N-phenylethylhydrazinecarbothioamide 4b in the form of light-yellow needle-shaped crystals. Yield: 0.94 g (86%).

^1^H NMR (400 MHz, CDCl_3_) δ 8.04 (br s, 1H), 7.48 (t, J = 5.8 Hz, 1H), 7.27 – 7.34 (m, 2H), 7.18 – 7.25 (m, 3H), 3.86 (q, J = 7.1 Hz, 2H), 3.73 (br s, 2H), 2.92 (t, J = 7.1 Hz, 2H).

**Synthesis of N-benzylhydrazinecarbothioamide (4c).**

From 1.35 g (0.009 mol) of benzyl isothiocyanate, as a result of the reaction with hydrazine hydrate, 0.931 g yielded N-benzylhydrazinecarbothioamide 4c in the form of white needle-shaped crystals. Yield: 1.34 g (81%).

^1^H NMR spectrum (400 MHz, CD_3_OD) δ 8.77 (br s, 1H,), 8.30 (br s, 1H), 7.30 -7.28 (m, 4H), 7.28-7.21 (m, 1H), 4.71 (d, 1H, J=6.2 Hz), 4.51 (br s, 2H).

**Synthesis of N-substituted 2-(3-oxobutan-2-yldiene) hydrazinecarbothioamides (5a-c).**

General technique:

N-substituted hydrazinecarbothioamides 4a-c (1 molar equivalent) were dissolved in MeOH (6.25 ml per 100 mg) and added dropwise to a solution of 2,3-butanedione (5 molar equivalents) in MeOH (4.5 ml per 100 µl) in the presence of catalytic quantities of HCl at 0°C for an hour. Then the reaction mixture was stirred at room temperature until the spot of N-substituted hydrazinecarbothioamides disappeared on the TLC of the reaction mixture. Then the solvent was evaporated under reduced pressure, and the resulting residue was purified using silica gel column chromatography in the system CH_2_Cl_2_:MeOH = 20:1.

**Synthesis of N-allyl-2-(3-oxobutan-2-yldiene) hydrazinecarbothioamide (5a).**

From 1 g of N-allylhydrazinecarbothioamide 4a, the reaction with 2,3-butanedione gave 3.28 g of N-allyl-2-(3-oxobutan-2-yldiene) hydrazinecarbothioamide 5a in the form of a white crystalline powder. Yield: 987 mg (65%).

^1^H NMR (400 MHz, CDCl_3_) δ 8.71 (s, 1H), 7.58 (s, 1H), 6.07 – 5.83 (m, 1H), 5.43 – 5.19 (m, 2H), 4.38 (t, J = 5.6 Hz, 2H), 2.41 (s, 3H), 2.01 (s, 3H).

^13^C NMR (100 MHz, CDCl_3_) δ 196.4, 178.3, 145.3, 132.7, 117.5, 47.0, 24.6, 9.2.

FTIR (Diamond, ν/cm^-1^): 621, 682, 834, 899, 912, 961, 1010, 1042, 1105, 1141, 1178, 1274, 1364, 1419, 1434, 1505, 1541, 1588, 1648, 1676, 2909, 2987, 3009, 3090, 3220, 3341.

HRMS C_8_H_13_N_3_OS (ESI TOF+MS) m/z: 200.0852 (Calc. for 200.0852 M+H^+^).

**Synthesis of N-phenylethyl-2-(3-oxobutan-2-yldiene)hydrazinecarbothioamide (5b).**

From 0.815 g of N-phenylethylhydrazinecarbothioamide 4b, the reaction with 2,3-butanedione gave 1.8 g of N-phenylethyl-2-(3-oxobutan-2-yldiene) hydrazinecarbothioamide 5b in the form of a white crystalline powder. Yield: 0.736 g (67%).

^1^H NMR (400 MHz, CD_3_OD) δ, 8.72 (s, 1H), 7.53 (s, 1H), 7.21 – 7.36 (m, 5H), 4.01 (q, J = 6.6 Hz, 2H), 3.00 (t, J = 6.7 Hz, 2H), 2.19 (s, 3H), 1.96 (s, 3H).

^13^C NMR (400 MHz, CDCl_3_) δ 196.4, 177.7, 144.9, 138.2, 129.0, 128.8, 127.0, 45.6, 34.8, 24.4, 8.9.

FTIR (Diamond, ν/cm^-1^): 701, 753, 996, 1130, 1175, 1209, 1232, 1298, 1360, 1428, 1454, 1503, 1538, 1608, 1688, 2937, 3026, 3174, 3337 (3113 16 02 23)

HRMS C_13_H_17_N_3_OS (ESI TOF+MS) m/z: 264.1164 (Calc. for 264.1165 M+H^+^).

**Synthesis of N-benzyl-2-(3-oxobutan-2-yldiene) hydrazinecarbothioamide (5c).**

From 1.5 g of N-benzylhydrazinecarbothioamide 4c, as a result of the reaction with 2,3-butanedione, 3.6 g gave N-benzyl-2-(3-oxobutan-2-yldiene) hydrazinecarbothioamide 5c in the form of a white crystalline powder. Yield: 1.2 g (59%).

^1^H NMR (400 MHz, CD_3_OD) δ 8.61 (br s, 1H), 8.21 (br s, 1H), 7.30-7.27 (m, 4H), 7.10-7.07 (m, 1H), 4.69 (d, 1H, J = 6.2 Hz), 2.11 (br s, 3H), 1.85 (br s, 3H).

**Synthesis of N-substituted 2-((Z)-3-(2-(5-(benzothiazol-2-yl)pyridin-2-yl)hydrazono)butan-2-ylidene)hydrazinecarbothioamides) (6a-c).**

General technique:

2-(6-hydrazinylpyridin-3-yl) benzothiazole 3 (1 molar equivalent) and N-substituted 2-(3-oxobutan-2-yldiene)hydrazinecarbothioamides 5a-c (1 molar equivalent) were stirred at boiling in EtOH (20 ml per 100 mg of compound 3) in a nitrogen atmosphere for 4 hours with the addition of catalytic quantities of AcOH. Then the reaction mixture was cooled, the precipitate that formed was filtered off, washed with EtOH, then with Et_2_O, and dried in air.

**Synthesis of N-allyl-2-((Z)-3-(2-(5-(benzothiazol-2-yl)pyridin-2-yl)hydrazono)butan-2-ylidene)hydrazinecarbothioamide) (6a).**

From 100 mg of N-allyl-2-(3-oxobutan-2-yldiene)-hydrazinecarbothioamide 5a, as a result of the reaction with 2-(6-hydrazinylpyridin-3-yl)benzothiazole 3, 121.6 mg was obtained N-allyl-2-((Z)-3-(2-(5-(benzothiazol-2-yl)pyridin-2-yl)hydrazono)butan-2-ylidene)-hydrazinecarbothioamide) 6a (87%) as light yellow powder.

^1^H NMR (400 MHz, CD_3_OD) δ 8.31 (dd, 1H, J_1_ = 8.78 Hz, J_2_ = 2.24 Hz), 8.03 (d, 1H, J = 7.94 Hz), 7.89 (d, 1H, J = 7.7 Hz), 7.49 (m, 1H), 7.43 (m, 2H), 5.97 (m, 1H), 5.30 (s, 1H), 5.24 (m, 1H), 4.38 (m, 2H,), 1.87 (s, 6H).

HRMS (M+H)^+^: calculated 424.13, found 424.15.

**Synthesis of N-phenylethyl-2-((Z)-3-(2-(5-(benzothiazol-2-yl)pyridin-2-yl)hydrazono)butan-2-ylidene)hydrazinecarbothioamide) (6b).**

From 100 mg of N-phenylethyl-2-(3-oxobutan-2-yldiene)hydrazinecarbothioamide 5b, as a result of the reaction with -(6-hydrazinylpyridin-3-yl)benzothiazole 3, 92 mg were obtained N-phenylethyl-2-((Z)- 3-(2-(5-(benzothiazol-2-yl)pyridin-2-yl)hydrazono)butan-2-ylidene) hydrazinecarbothioamide) 6b in the form of a yellow powder.

^1^H NMR (400 MHz, CDCl_3_) δ 8.89 (s, 1H), 8.65 (s, 1H), 8.42 (s, 1H), 8.32 (d, J = 8.80 Hz, 1H), 8.05 (d, J = 7.83 Hz, 1H), 7.91 (d, J = 7.82 Hz, 1H), 7.5 (m, 2H), 7.32 – 7.24 (m, 4H), 4.02 (q, J = 5.87 Hz, 2H), 3.01 (q, J = 5.87 Hz, 2H), 2.22 (s, 3H), 1.97 (s, 3H).

(M+H)^+^: calculated 488.16, found 488.20.

**Synthesis of N-benzyl-2-((Z)-3-(2-(5-(benzothiazol-2-yl)pyridin-2-yl)hydrazono)butan-2-ylidene)hydrazinecarbothioamide) (6c).**

From 100 mg of N-benzyl-2-(3-oxobutan-2-yldiene)hydrazinecarbothioamide 5c, as a result of the reaction with 2-(6-hydrazinylpyridin-3-yl)benzothiazole 3, 97.2 mg were obtained N-benzyl-2-((Z)-3-(2-(5-(benzothiazol-2-yl)pyridin-2-yl)hydrazono)butan-2-ylidene)hydrazinecarbothioamide) 6c (67%) as a yellow powder.

^1^H NMR (400 MHz, CDCl_3_) δ 9.24 (s, 1H), 9.11 (s, 1H), 8.76 (s, 1H, Ar), 8.18 (d, J = 8.80 Hz, 1H), 7.91 (m, 2H), 7.81 (m, 1H), 7.81 (d, J = 7.82 Hz, 2H), 7.43 – 7.21 (m, 4H), 4.88 (s, 2H), 3.01 (q, J = 5.87 Hz, 2H), 2.22 (s, 3H), 2.05 (s, 3H).

HRMS (M+H)^+^: calculated 474.14, found 474.15.

**Synthesis of carboxymethyl-N-methyl-N-phenyldithiocarbamate (7).**

A mixture of carbon disulfide (12 ml, 0.20 mol) and N-methylaniline (21.6 ml, 0.20 mol) was treated with sodium hydroxide (8.2 g, 0.20 mol) dissolved in 250 ml of water. The organic layer disappears after stirring the solution for 4 hours at room temperature. At this point, the resulting straw-colored solution was treated with sodium chloroacetate (23.2 g, 0.20 mol) and left for 24 hours. The resulting solution was acidified with concentrated HCl solution and solids were separated, collected and dried. 38.56 g (80%) of light-yellow powder was obtained. Tpl. = 199 – 200 °C. (197 – 198 °С lit).

^1^H NMR (400 MHz, DMSO-d6) δ 12.69 (br s, 1H), 7.39-7.53 (m, 5H), 3.96 (s, 2H), 3.67 (s, 3H).

**Synthesis of 4-methyl-4-phenyl-3-thiosemicarbazide (8).**

A solution of 17.7 g (0.0733 mol) of carboxymethyl-N-methyl-N-phenyldithiocarbamate 7 in 20 ml of 98% hydrazine hydrate and 10 ml of water was heated in a water bath (85°C) until crystals separated. The crystals were filtered, washed with water, and dried in a vacuum. The crude product was recrystallized from a mixture of 50 ml EtOH and 25 ml water. 9.1 g (68%) of thick, colorless rods of 4-methyl-4-phenyl-3-thiosemicarbazide were obtained. T_melt_ = 115 – 116 °С (112 – 113^o^C lit.).

^1^H NMR (400 MHz, DMSO-*d6*) δ 8.60 (br s, 1H), 7.10 – 7.49 (m, 5H), 4.80 (br.s, 2H), 3.42 (s, 3H).

^13^C NMR (100 MHz, CDCl_3_) δ 181.9, 138.8, 128.8, 128.7, 126.6, 45.1, 35.5.

**Synthesis of (E)-2-((E)-3-(2-(allylcarbamothioyl)hydrazono)butan-2-ylidene)-N-methyl-N-phenylhydrazinecarbothioamide (9a).**

In a round bottom flask equipped with a reflux condenser, a mixture of diacetyl-mono-4-N-phenylethyl-3-thiosemicarbazone 5a (1.46 g, 5.6 mmol) and 4-methyl-4-phenyl-3-thiosemicarbazide 8 (1 g, 5 .6 mmol) was boiled under argon in dry methanol (20 ml) in the presence of catalytic quantities of acetic acid. The reaction was carried out for 6 hours until a precipitate formed. The reaction mixture was cooled to room temperature, the precipitate was filtered off, and washed with a small amount of methanol. The result was 1.43 g (63%) of product. T_melt_ = 78 – 79^o^C.

^1^H NMR (400 MHz, DMSO-*d6*) δ 10.88 (s, 1H), 10.27-10.45 (m, 2H), 9.32 (d, J = 4.2 Hz, 1H), 8.56 (t, J = 6.0 Hz, 1H), 7.33-7.57 (m, 5H), 5.83-5.98 (m, 1H), 5.04-5.20 (m, 2H), 4.23 (t, J = 5.7 Hz, 2H), 3.56 (s, 3H), 2.22 (s, 3H), 2.19 (s, 3H).

^13^C NMR (100 MHz, DMSO-*d6*) δ 179.7, 178.1, 176.9, 175.6, 149.5, 147.6, 143.1, 134.8, 130.0, 127.9, 126.7, 115.6, 46.0, 43.0, 12.0, 11.6.

FTIR (Diamond, ν/cm^-1^): 619, 669, 696, 742, 770, 808, 851, 874, 908, 932, 954, 1055, 1106, 1130, 1207, 1238, 1264, 1290, 1306, 1359, 1414, 1462, 1529, 1585, 1594, 1649, 2912, 2977, 3184, 3346.

**Synthesis of (E)-2-((E)-3-(2-(phenylethylcarbamothioyl)hydrazono)butan-2-ylidene)-N-methyl-N-phenylhydrazinecarbothioamide (9b).**

In a round bottom flask equipped with a reflux condenser, a mixture of diacetyl-mono-4-N-phenylethyl-3-thiosemicarbazone 5b (1.46 g, 5.6 mmol) and 4-methyl-4-phenyl-3-thiosemicarbazide 8 (1 g, 5.6 mmol) was boiled under argon in dry methanol (20 ml) in the presence of catalytic amounts of acetic acid. The reaction was carried out for 6 hours until a precipitate formed. The reaction mixture was cooled to room temperature, the precipitate was filtered off, and washed with a small amount of methanol. The result was 1.43 g (60%) of product.

^1^H NMR (400 MHz, DMSO-d6) δ 10.89 (s, 1H), 10.40 (s, 1H), 10.35 (br s, 1H), 9.32 (br s, 1H), 8.40 (t, J = 5.9 Hz, 1H), 7.14-7.56 (m, 11H), 3.73-3.86 (m, 2H), 3.56 (s, 3H), 2.91 (t, J = 7.6 Hz, 3H), 2.19 (s, 3H), 2.17 (s, 3H).

^13^C NMR (100 MHz, DMSO-d6) δ 206.6, 179.7, 177.7, 175.5, 149.3, 147.4, 143.0, 139.1, 129.9, 128.6, 128.4, 127.9, 126.7, 126.2, 45.2, 43.0, 34.5, 11.9, 11.6.

FTIR (Diamond, ν/cm-1): 3338, 3218, 2946, 1595, 1529, 1490, 1471, 1409, 1366, 1267, 1208, 1133, 1065, 698.

**Synthesis of 4-bromomethylnitrobenzene (10).**

To a solution of 4-nitrotoluene (72.9 mmol, 1 eq.) in carbon tetrachloride (100 ml), azobiisobutyronitrile (AIBN) (3.7 mmol, 0.05 eq.) was added as a radical generator. N-bromosuccinimide (NBS) (76.6 mmol, 1.05 eq.) was added to this suspension and the mixture was stirred at reflux for 6 hours. The reaction mixture was filtered and concentrated to give a dull yellow solid, which was recrystallized from hexane several times, yield: 10.70 g (68%).

^1^H NMR (CDCl_3_, 400 MHz) δ 8.20 (d, 2H, J = 8.6 Hz), 7.57 (d, 2H, J = 8.6 Hz), 4.52 (s, 2H).

**Synthesis of (E)-4-nitro-4'-methylaminostilbene (11).**

4-bromomethylnitrobenzene 10 (1.51 g, 7.0 mmol) was dissolved in 5 ml of P(OEt)_3_ and was heated for 2 hours at reflux. Then reaction mixture was cooled to room temperature and excess of triethyl phosphite was removed under reduced pressure. Obtained oil was dissolved in dimethylformamide (10 ml) and then sodium hydride (60% in mineral oil, 0.39 g, 9.8 mmol) was added. The reaction was stirred for 15 min followed by the addition of 4-methylaminobenzaldehyde (0.94 g, 7.0 mmol). The mixture was stirred for 2 hours, and then ethanol (2 ml), water (50 ml) was added until a red precipitate formed. The precipitate is separated by filtration and washed with water. The red solid was dissolved in dichloromethane and washed with H_2_O (3 x 200 ml) and then brine (1 x 100 ml). The organic phase was dried over MgSO4) and filtered, the solvent was removed under reduced pressure to give a red solid (1.4 g, 5.5 mmol, 75%).

^1^H NMR (400 MHz, CDCl_3_) δ 8.18 (m, 2H), 7.56 (m, 2H), 7.45 (m, 2H), 7.21 (d, J = 16.2 Hz, 1H), 6.93 (d, J = 16.2 Hz, 1H), 6.72 (m, 2H), 3.02 (s, 6H).

**Synthesis of (E)-4-amino-4'-dimethylaminostilbene (12).**

(E)-4-nitro-4'-dimethylaminostilbene 11 (1.7 g, 6.5 mmol) was dissolved in 40 ml ethanol and the mixture was bubbled with Ar for 1 hour. Stannous chloride (7.6 g, 40 mmol) was then added to this mixture and refluxed. After 1.5 hours, the reaction mixture was cooled to ambient room temperature and adjusted to pH 9 (1 M NaOH). Then extraction was carried out with dichloromethane (300 ml), the organic phase was washed with brine (150 ml), dried over MgSO_4_); the solvent was then removed under reduced pressure to give a yellow solid (1.1 g, 4.6 mmol, 68%). T_melt_ = 148-149˚С (172 – 173^o^C lit).

^1^H NMR (400 MHz, DMSO-*d6*) δ 7.32 (d, J = 8.9 Hz, 2H), 7.20 (d, J = 8.5 Hz, 2H), 6.78 (s, 2H), 6.68 (d, J = 8.9 Hz, 2H), 6.54 (d, J = 8.5 Hz, 2H), 2.89 (s, 6H).

**Synthesis of (E)-N-allyl-2-((E)-3-(2-((4-((E)-4-(dimethylamino)styryl)phenyl)carbamothioyl) hydrazono)butan-2-ylidene)hydrazinecarbothioamide (13a).**

In a round bottom flask equipped with a reflux condenser, a mixture of N-phenyl-N-methyl-N'-allyl bis(acetylthiosemicarbazone) 9a (1.21 g, 3.32 mmol) and (E)-4-amino-4’-dimethylaminostilbene 12 (0.792 g, 3.32 mmol) was refluxed under argon in dry acetonitrile (30 ml) in the presence of catalytic amounts of acetic acid. The reaction was carried out for 20 hours. The reaction mixture was cooled to room temperature, the precipitate was filtered off, and washed with a small amount of acetonitrile. As a result, 1.06 g of product 13a (60%) was obtained.

^1^H NMR (400 MHz, DMSO*-d6*) δ 10.60 (s, 1H), 10.37 (s, 1H), 9.96 (s, 1H), 8.58 (t, 1H), 7.49-7.76 (m, 4H), 7.43 (d, J = 8.4 Hz, 3H), 7.11 (d, J = 16.4 Hz, 1H), 6.97 (d, J = 16.4, 1H), 6.72 (d, J = 8.4 Hz, 4H), 5.81-6.05 (m, 1H), 5.13 (dd, J_1_ = 13.5 Hz, J_2_ = 19.6 Hz, 2H), 2.93 (s, 6H), 2.29 (s, 3H), 2.27 (s, 3H)z, Hz1H.

^13^C NMR (100 MHz, DMSO*-d6*) δ 261.7, 178.1, 176.5, 150.0, 149.2, 148.1, 137.4, 135.1, 134.8, 128.4, 127.5, 125.4, 125.3, 125.0, 123.0, 115.6, 112.2, 46.0, 40.0, 12.1, 11.8.

FTIR (Diamond, ν/cm^-1^): 625, 644, 678, 707, 721, 742, 783, 802, 822, 862, 928, 950, 966, 1012, 1059, 1130, 1166, 1191, 1201, 1249, 1321, 1351, 1412, 1485, 1518, 1580, 1605, 2789, 2839, 2879, 2974, 3017, 3194, 3289, 3364.

HRMS: C_25_H_31_N_7_S_2_ Calculated: 493.2082; found 493.1698.

**Synthesis of (E)-N-phenylethyl-2-((E)-3-(2-((4-((E)-4-(dimethylamino)styryl)phenyl) carbamothioyl)hydrazono)butan-2-ylidene)hydrazinecarbothioamide (13b).**

In a round bottom flask equipped with a reflux condenser, a mixture of N-phenyl-N-methyl-N'-phenylethyl bis(acetylthiosemicarbazone) (30 mg, 81 mmol) 9b and (E)-4-amino-4’-dimethylaminostilbene 12 (20 mg, 81 mmol) was boiled under argon in dry acetonitrile (10 ml) in the presence of catalytic quantities of acetic acid. The reaction was carried out for 20 hours. The reaction mixture was cooled to room temperature, the precipitate was filtered off, and washed with a small amount of acetonitrile. The result was 27 mg of product (60%)

^1^H NMR (400 MHz, DMSO-*d6*) δ 10.60 (s, 1H), 10.39 (s, 1H), 9.95 (s, 1H), 8.42 (t, J = 5.7 Hz, 1H), 7.47-7.60 (m, 4H), 7.42 (d, J = 8.4 Hz, 2H), 7.26 (m, 5H), 7.11 (d, J = 16.3 Hz, 1H), 6.96 (d, J = 16.4 Hz, 1H), 6.71, d, J = 8.5 Hz, 2H), 3.78 (q, J = 6.7 Hz, 2H), 2.92 (s, 8H), 2.25 (s, 3H), 2.22 (s, 3H).

^13^C NMR (100 MHz, DMSO-*d6*) δ 177.7, 176.4, 149.9, 149.0, 147.9, 139.1, 137.4, 135.0, 128.6, 128.5, 128.4, 127.4, 126.2, 125.4, 125.0, 123.0, 112.2, 45.2, 40.0, 34.5, 12.0, 11.8.

FTIR (Diamond, ν/cm^-1^): 825, 967, 1078, 1135, 1185, 1253, 1343, 1419, 1490, 1525, 1582, 1607, 2792, 2847, 2881, 2935, 3021, 3222, 3295, 3368.

### NMR and FTIR spectra & LC-MS and HR-MS data

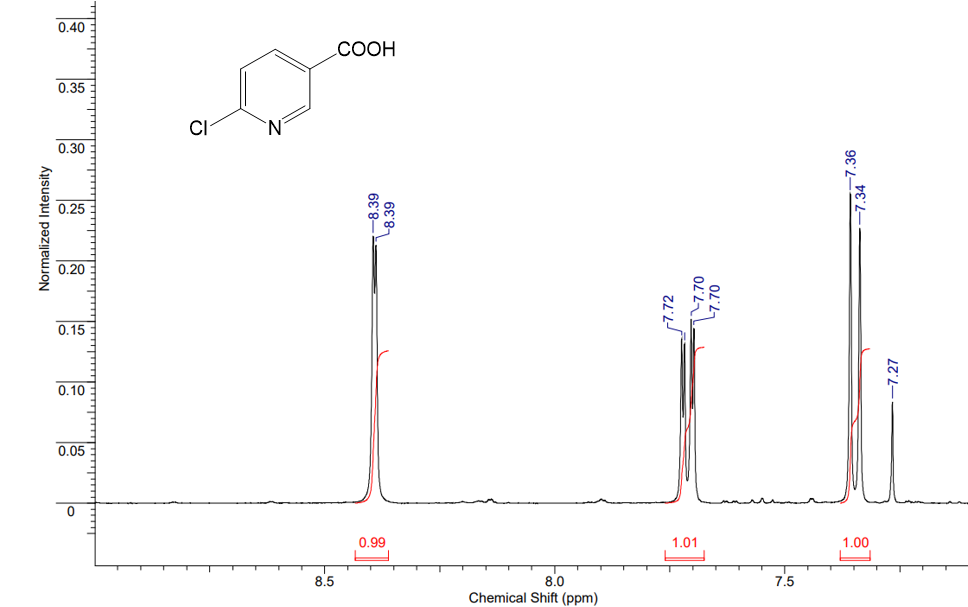

Figure S1. NMR ^1^H spectrum of 6-chloronicotinyl chloride (1)

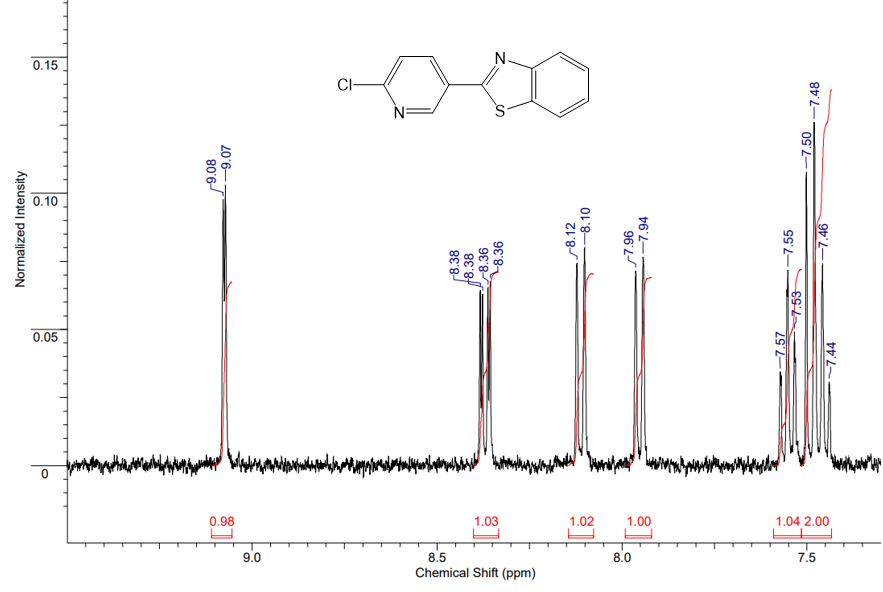

Figure S2. NMR ^1^H spectrum of 2-(6-chloropyridin-3-yl) benzothiazole (2)

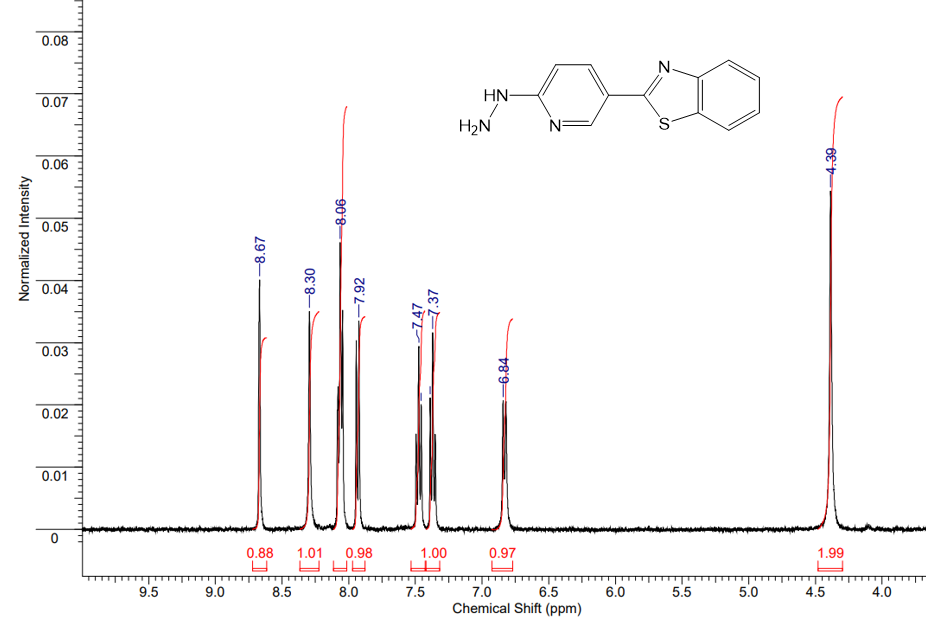

Figure S3. NMR ^1^H spectrum of 2-(6-hydrazinylpyridin-3-yl) benzothiazole (3)

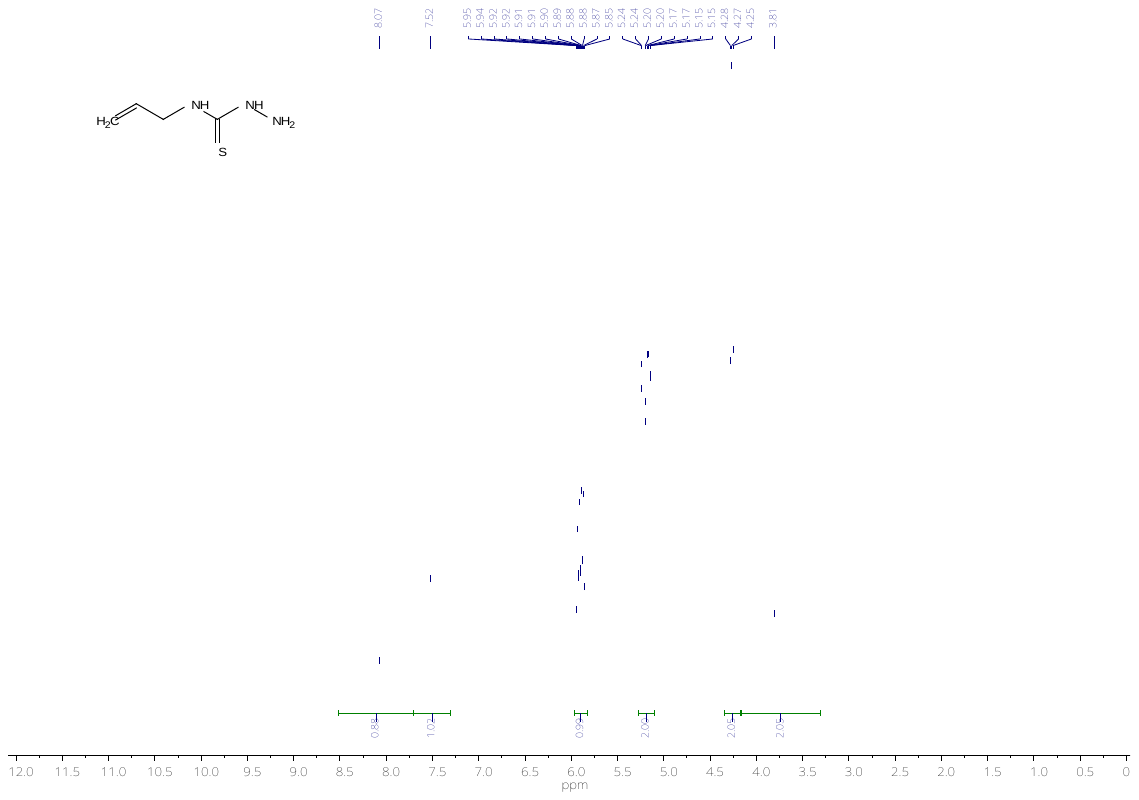

Figure S4. NMR ^1^H spectrum of N-allylhydrazinecarbothioamide (4a)

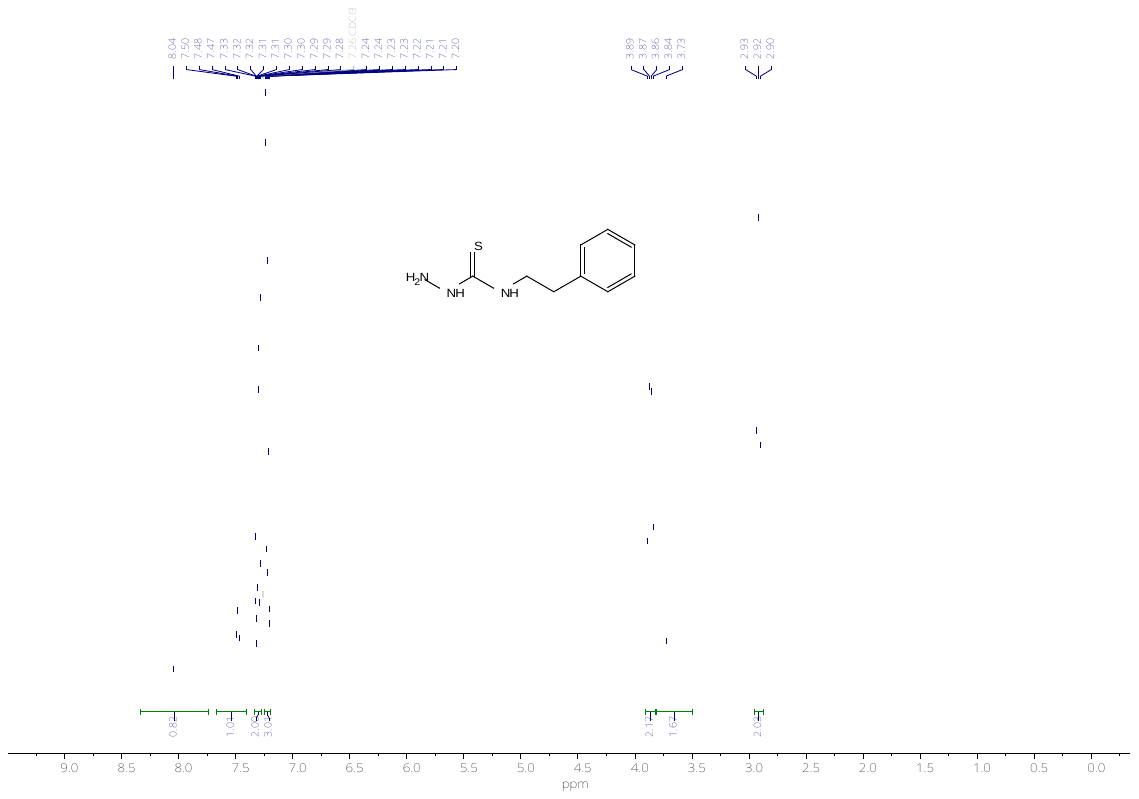

Figure S5. NMR ^1^H spectrum of phenylethylhydrazinecarbothioamide (4b)

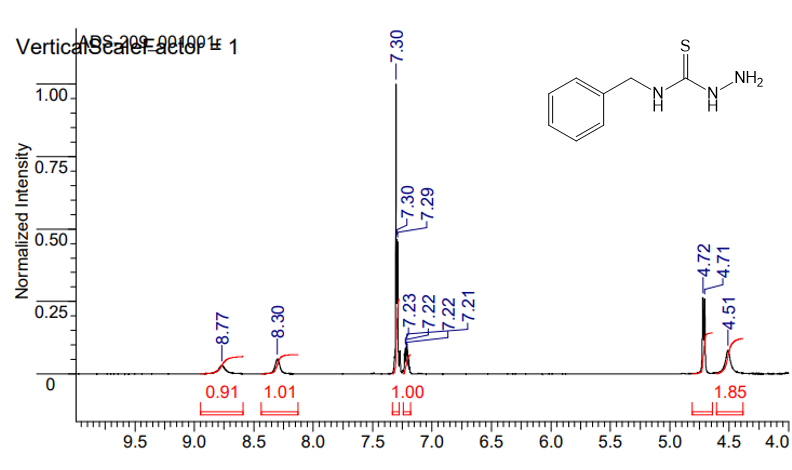

Figure S6. NMR ^1^H spectrum of benzylhydrazinecarbothioamide (4c)

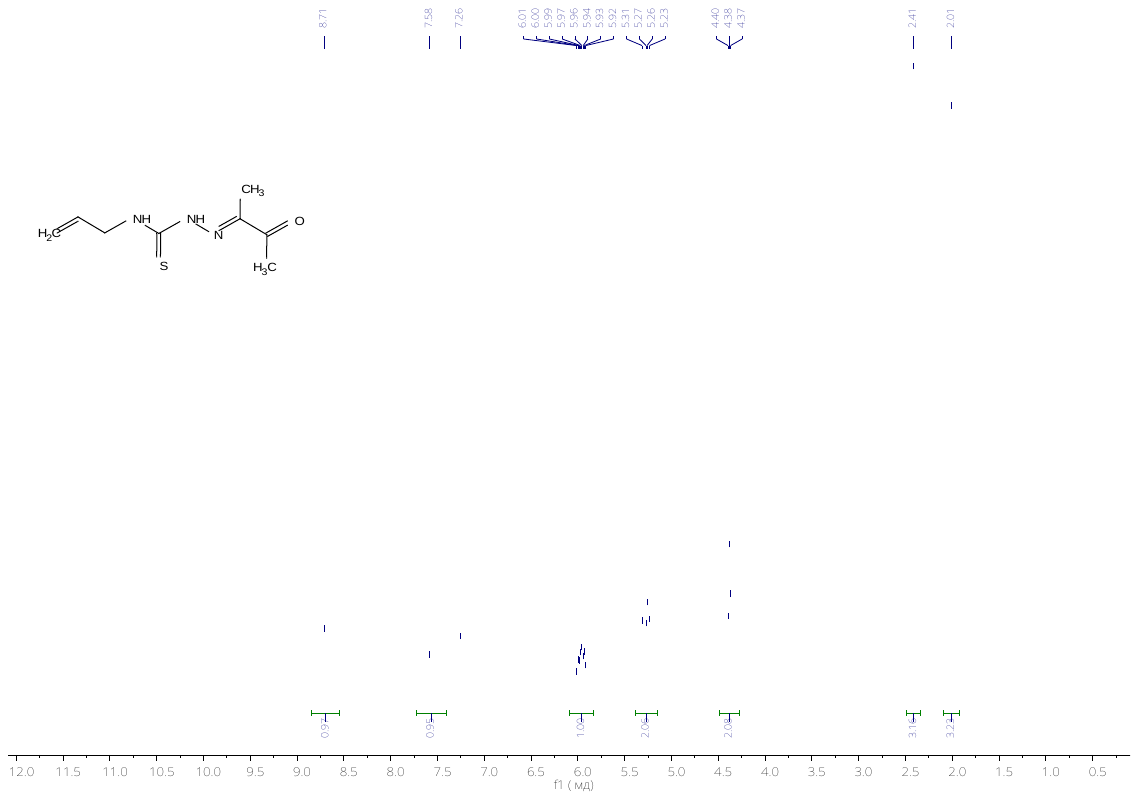

Figure S7. NMR ^1^H N-allyl-2-(3-oxobutan-2-yldiene) hydrazinecarbothioamide (5a)

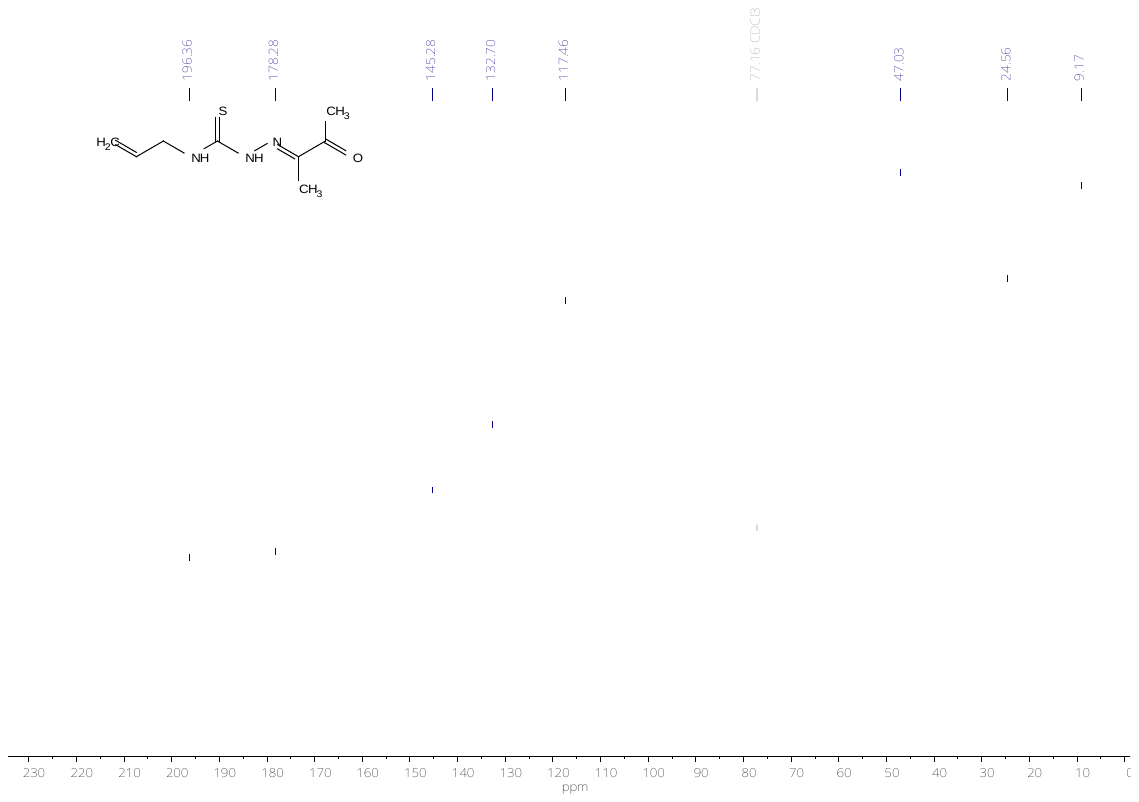

Figure S8. NMR ^13^C spectrum of N-allyl-2-(3-oxobutan-2-yldiene) hydrazinecarbothioamide (5a)

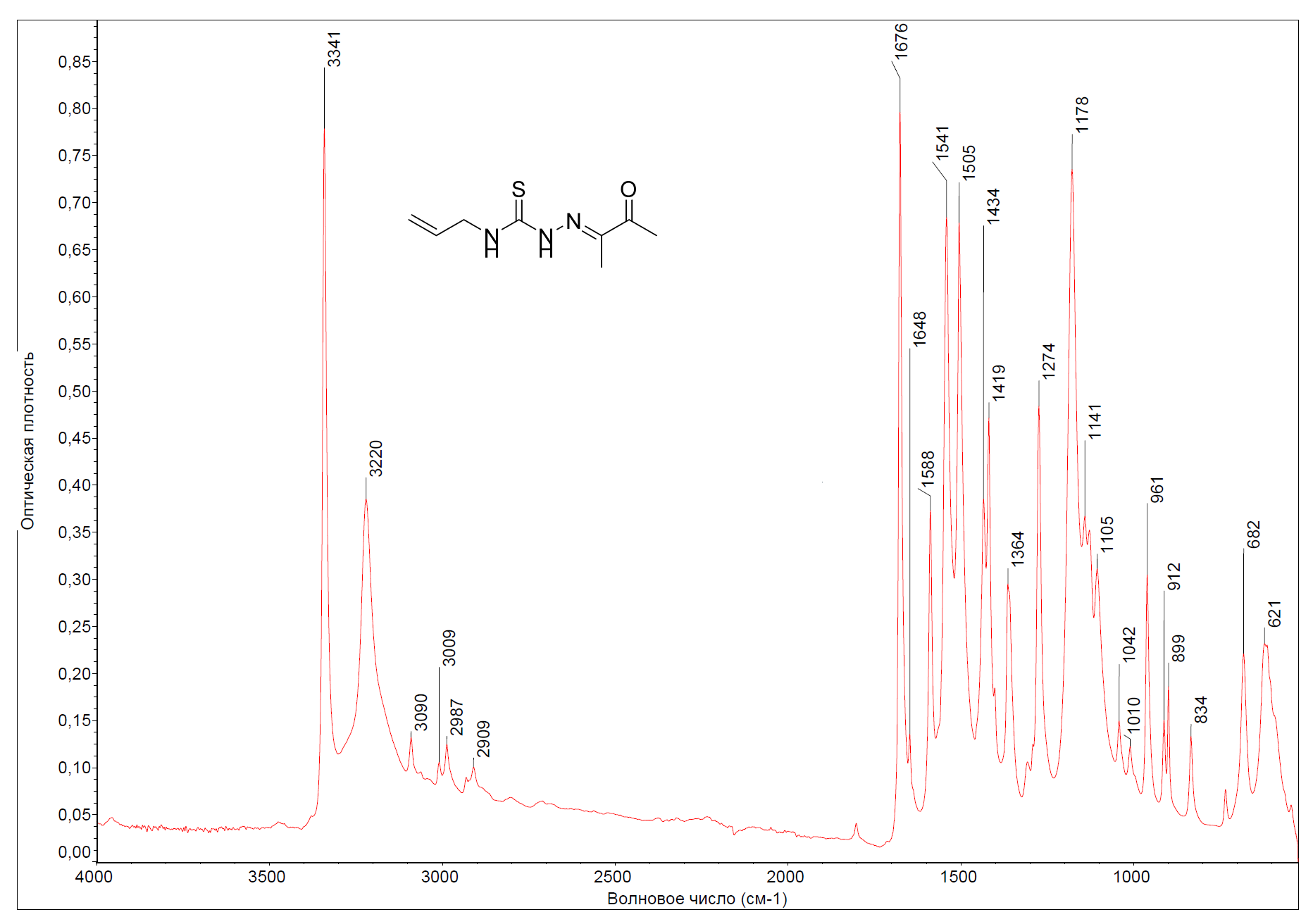

Figure S9. FTIR spectrum of N-allyl-2-(3-oxobutan-2-yldiene) hydrazinecarbothioamide (5a)

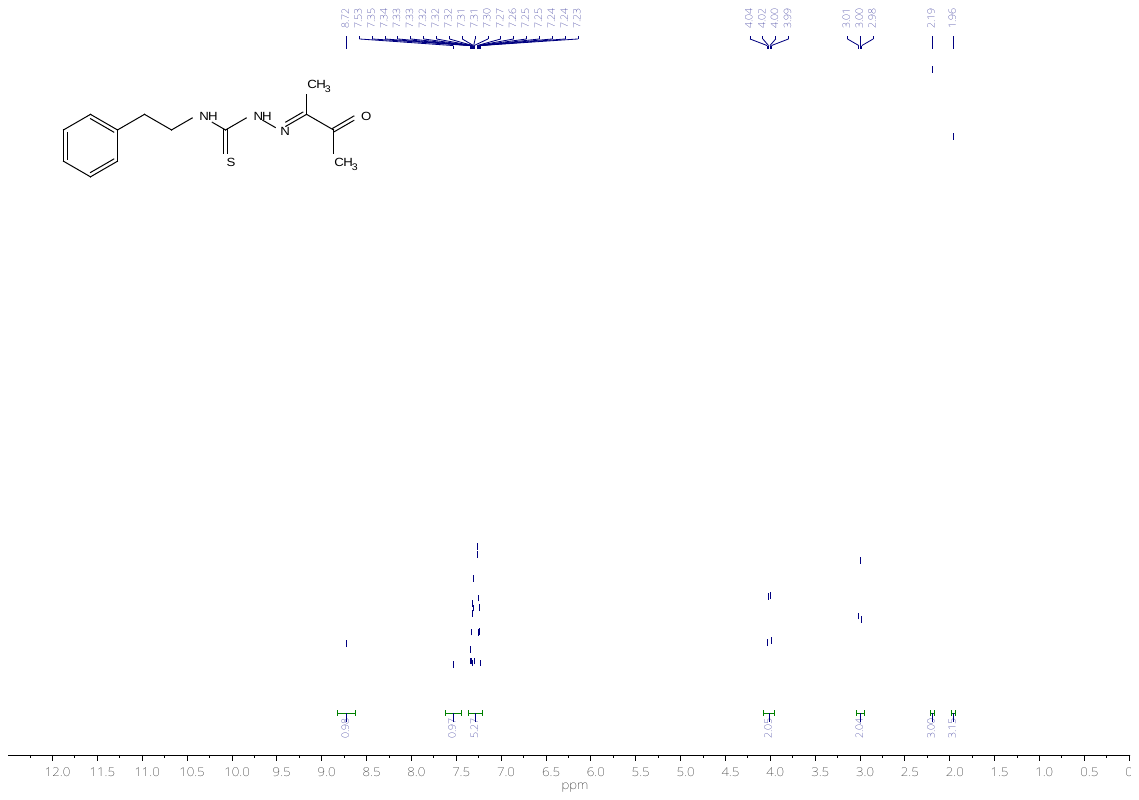

Figure S10. NMR ^1^H spectrum of N-phenylethyl-2-(3-oxobutan-2-yldiene)hydrazinecarbothioamide (5b)

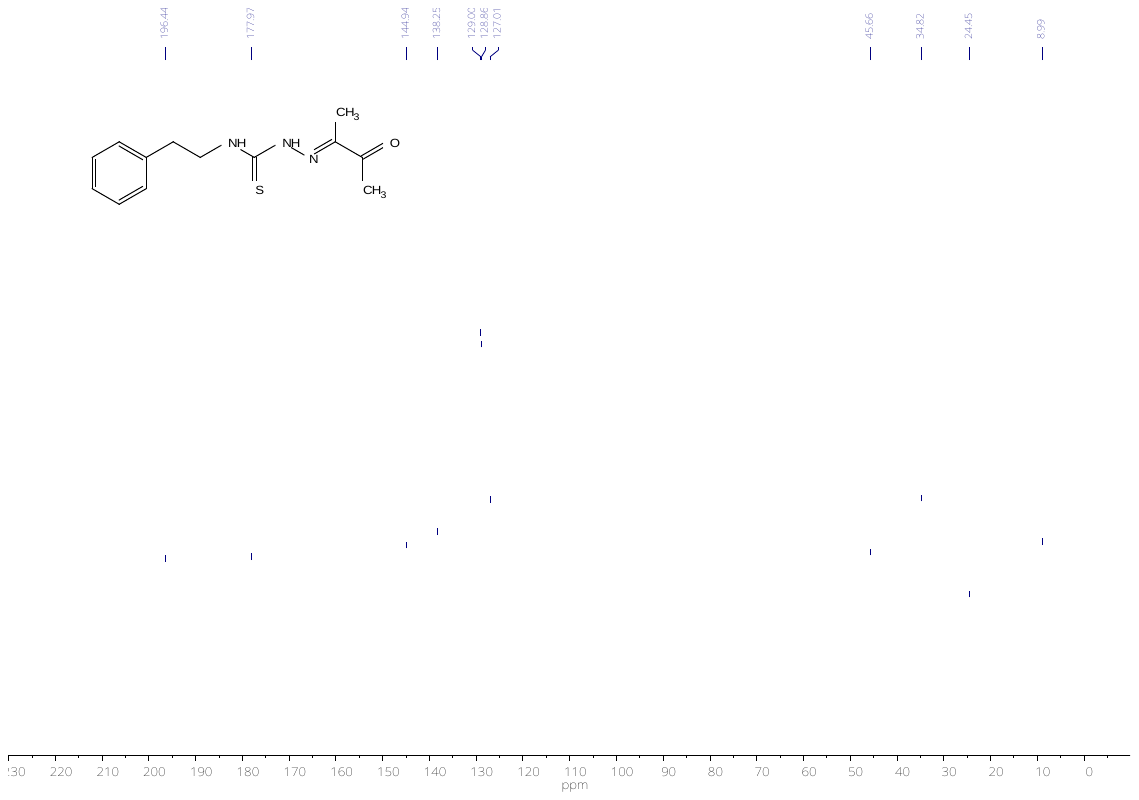

Figure S11. NMR ^13^C spectrum of N-phenylethyl-2-(3-oxobutan-2-yldiene)hydrazinecarbothioamide (5b)

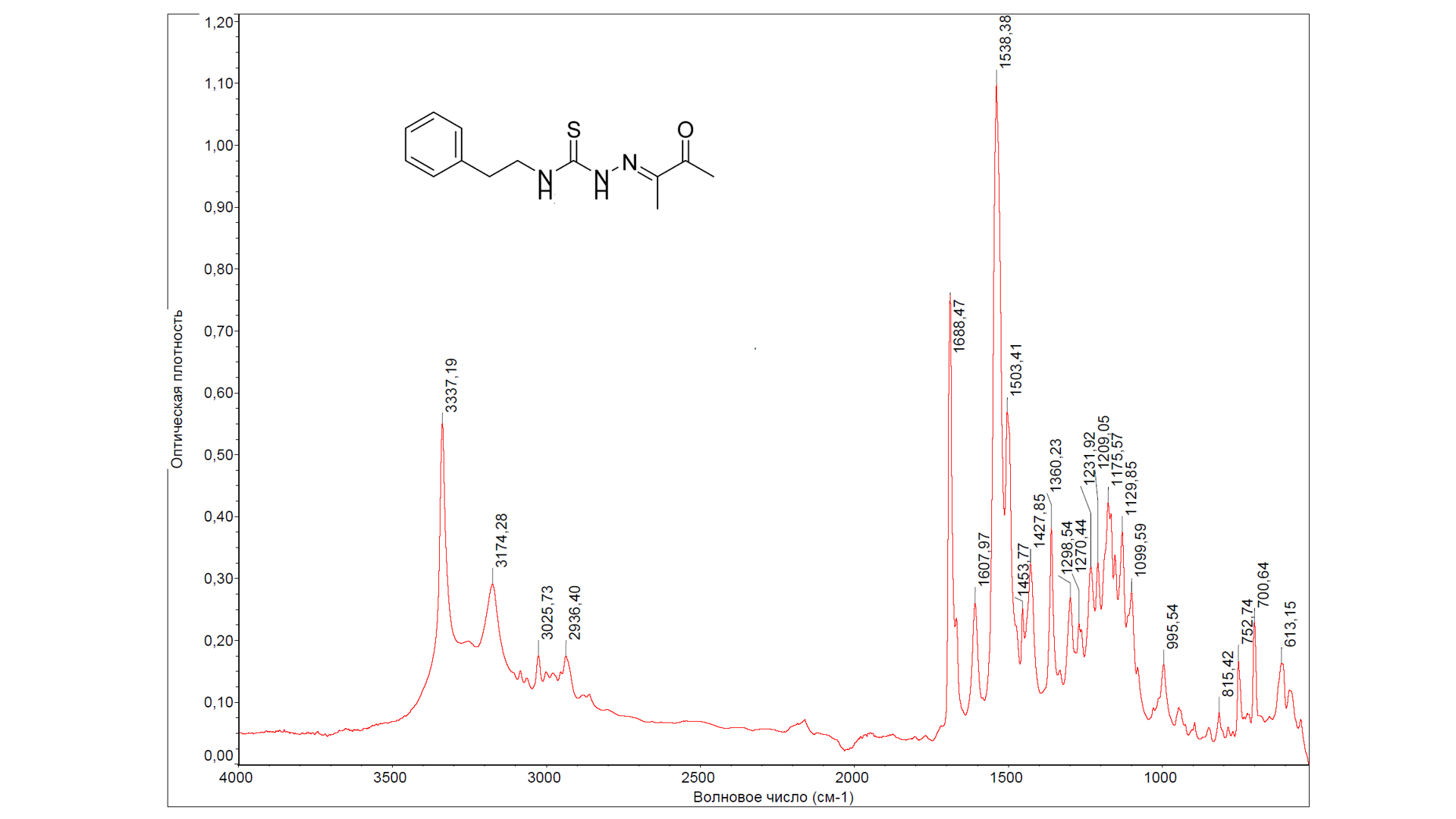

Figure S12. FTIR spectrum of N-phenylethyl-2-(3-oxobutan-2-yldiene)hydrazinecarbothioamide (5b)

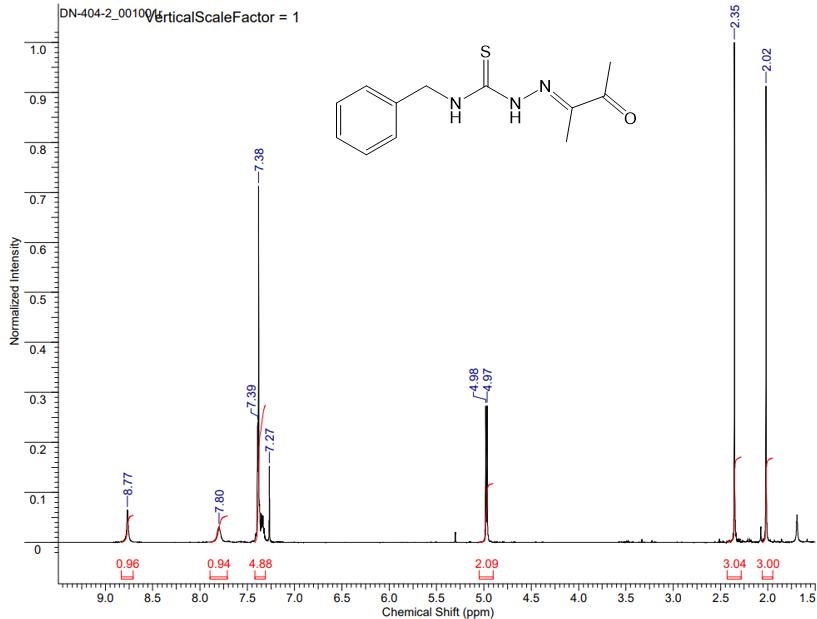

Figure S13. NMR^1^H spectrum of N-benzyl-2-(3-oxobutan-2-yldiene) hydrazinecarbothioamide (5с)

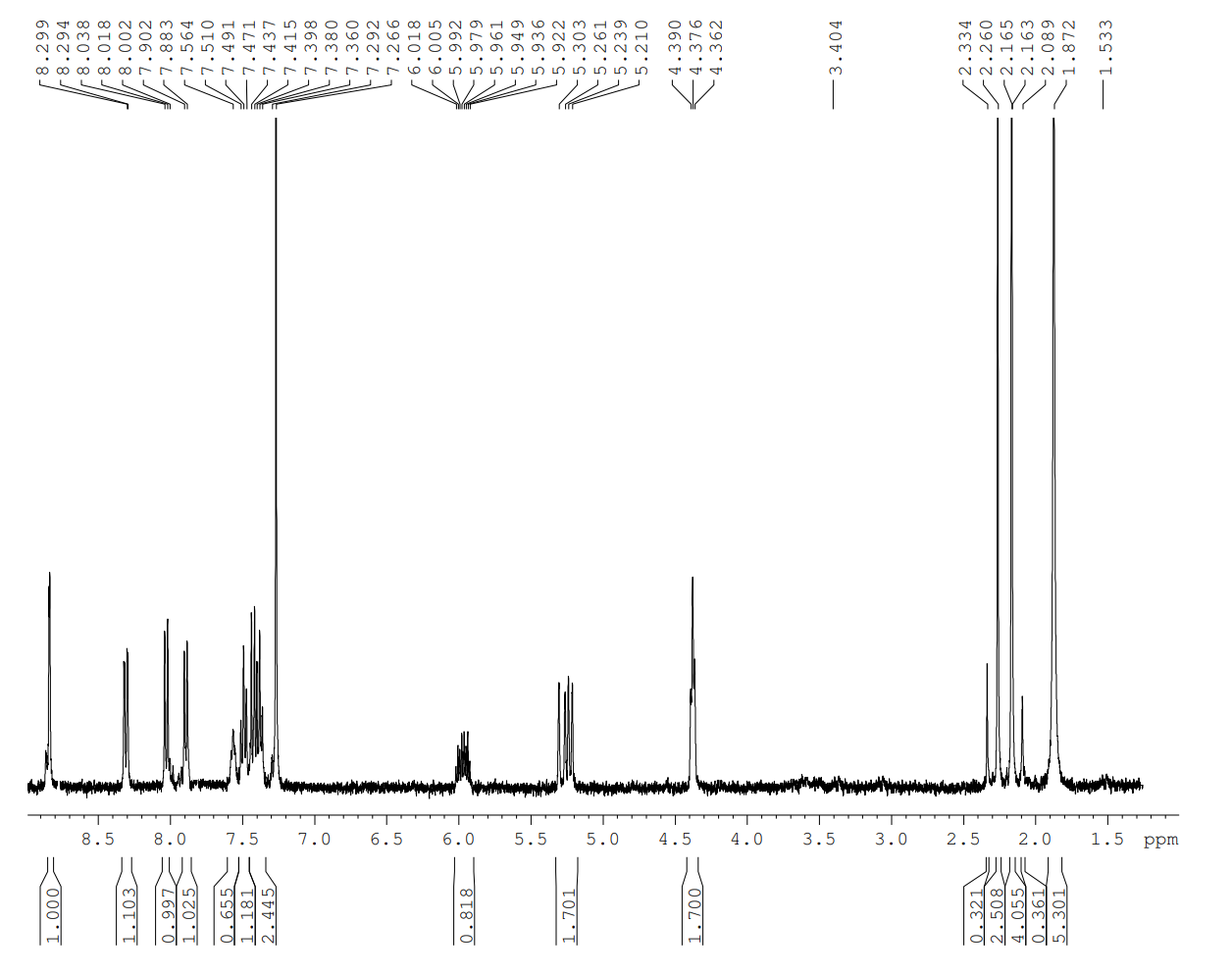

Figure S14. NMR ^1^H spectrum of N-allyl-2-((Z)-3-(2-(5-(benzothiazol-2-yl) pyridin-2-yl)hydrazono)butan-2-ylidene)hydrazinecarbothioamide) (6a)

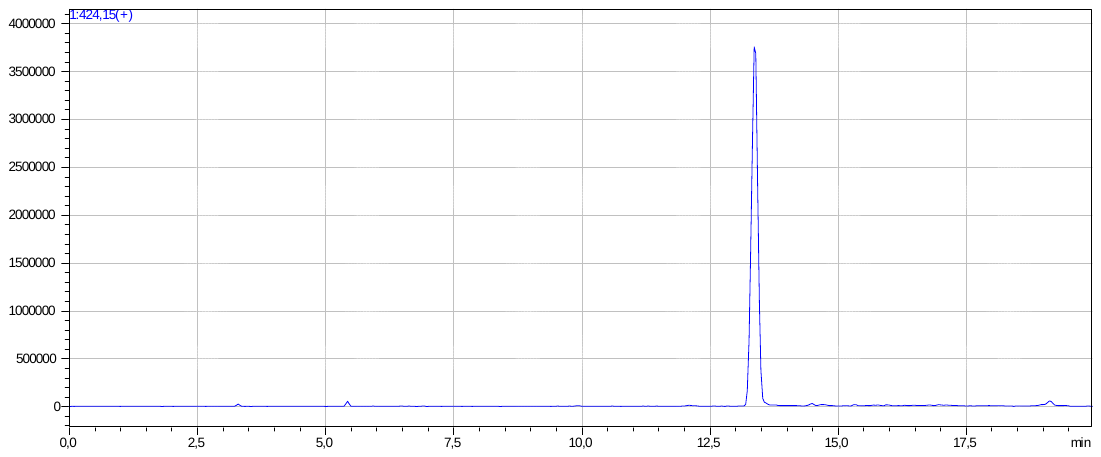

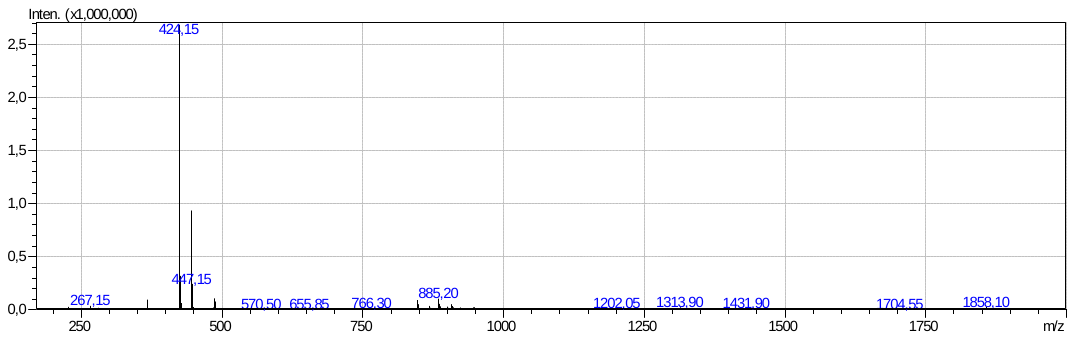

Figure S15. LC-MS data on N-allyl-2-((Z)-3-(2-(5-(benzothiazol-2-yl)pyridin-2-yl)hydrazono)butan-2-ylidene)hydrazinecarbothioamide) (6a)

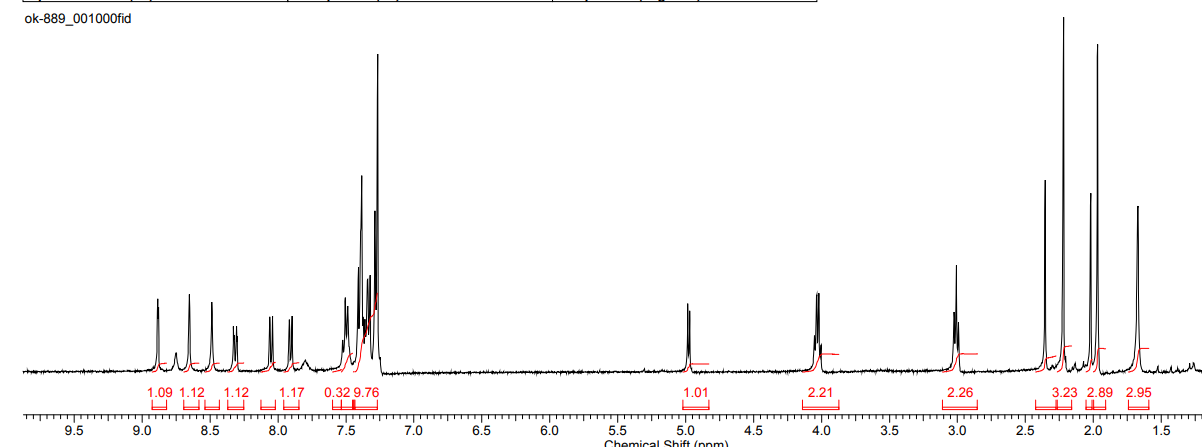

Figure S16. NMR ^1^H spectrum of N-phenylethyl-2-((Z)-3-(2-(5-(benzothiazol-2-yl)pyridin-2-yl)hydrazono)butan-2-ylidene)hydrazinecarbothioamide) (6b)

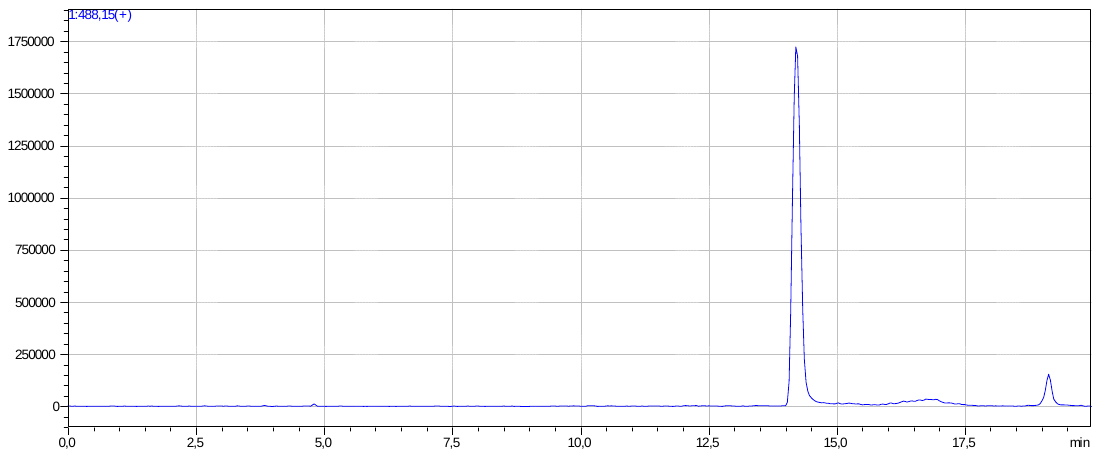

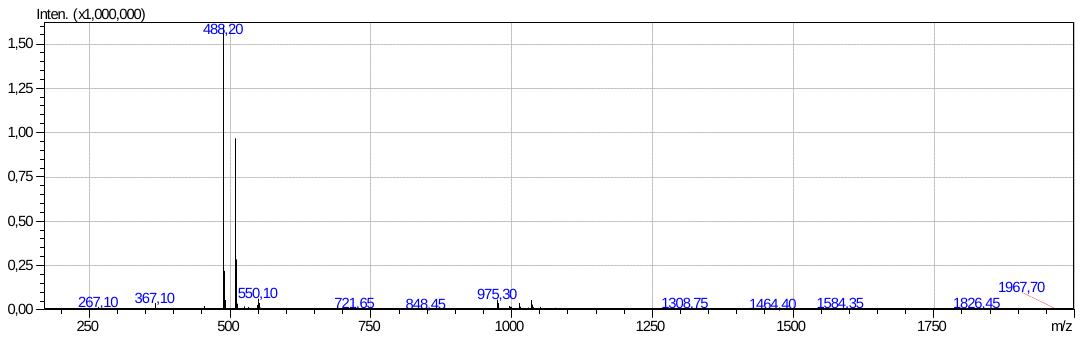

Figure S17. LC-MS data on N-phenylethyl-2-((Z)-3-(2-(5-(benzothiazol-2-yl)pyridin-2-yl)hydrazono) butan-2-ylidene)hydrazinecarbothioamide) (6b)

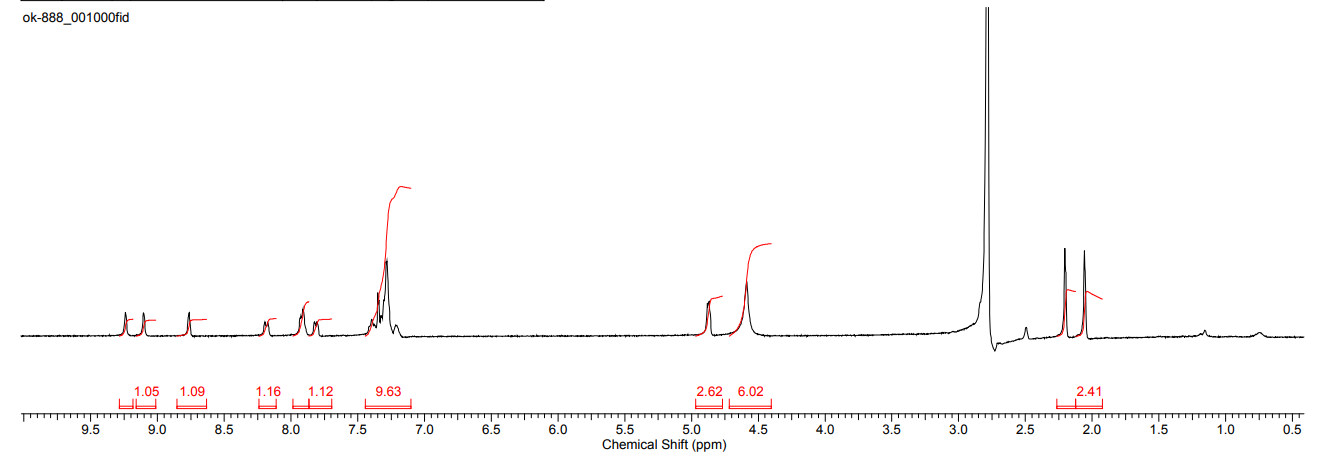

Figure S18. NMR ^1^H spectrum of N-benzyl-2-((Z)-3-(2-(5-(benzothiazol-2-yl)pyridin-2-yl)hydrazono)butan-2-ylidene)hydrazinecarbothioamide) (6c)

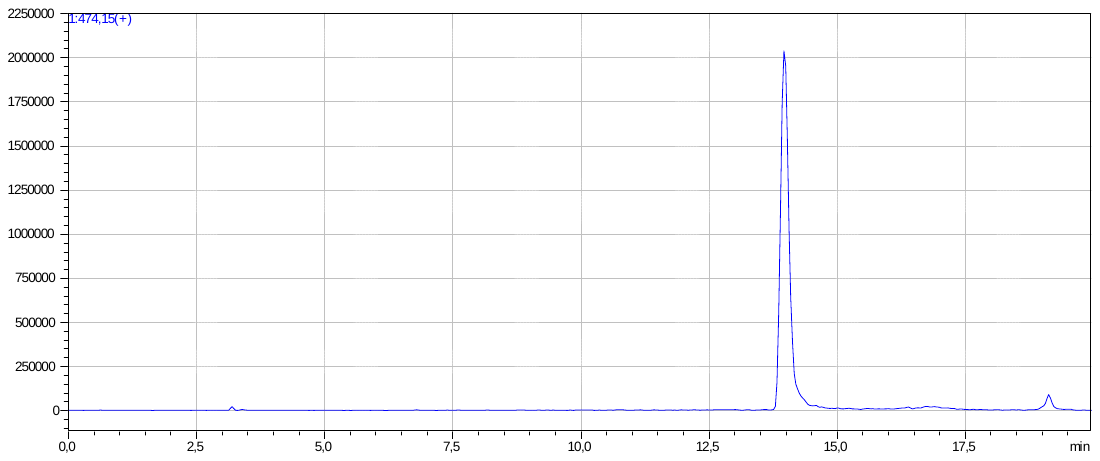

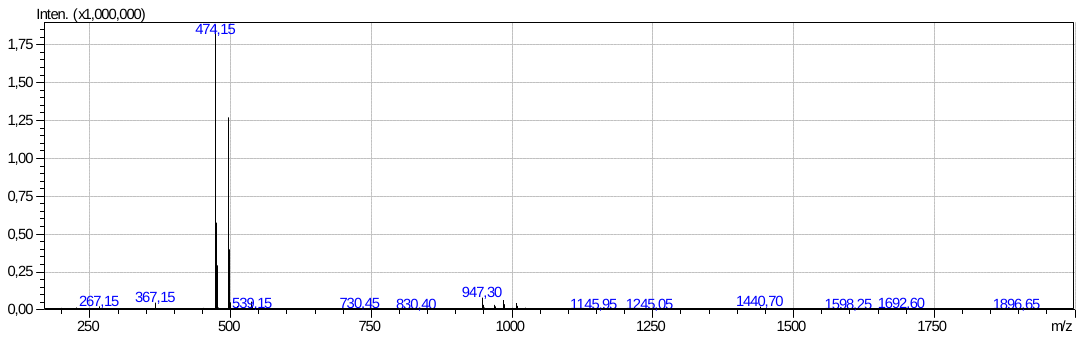

Figure S19. LC-MS data on N-benzyl-2-((Z)-3-(2-(5-(benzothiazol-2-yl)pyridin-2-yl)hydrazono)butan-2-ylidene)hydrazinecarbothioamide) (6c)

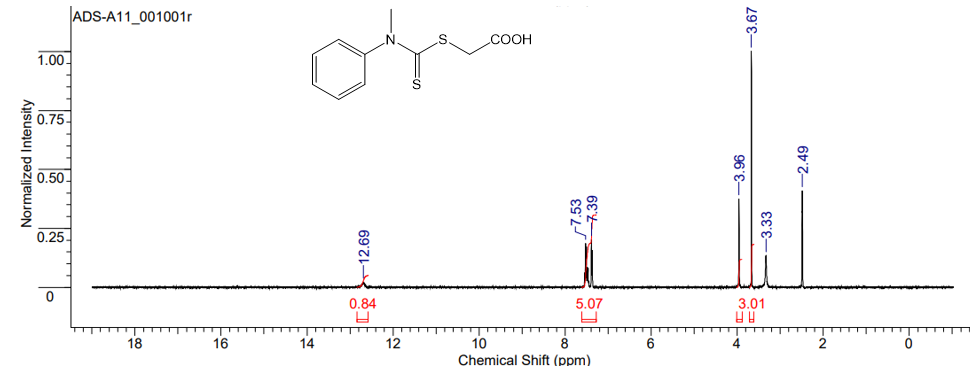

Figure S20. NMR ^1^H spectrum of carboxymethyl-N-methyl-N-phenyldithiocarbamate (7)

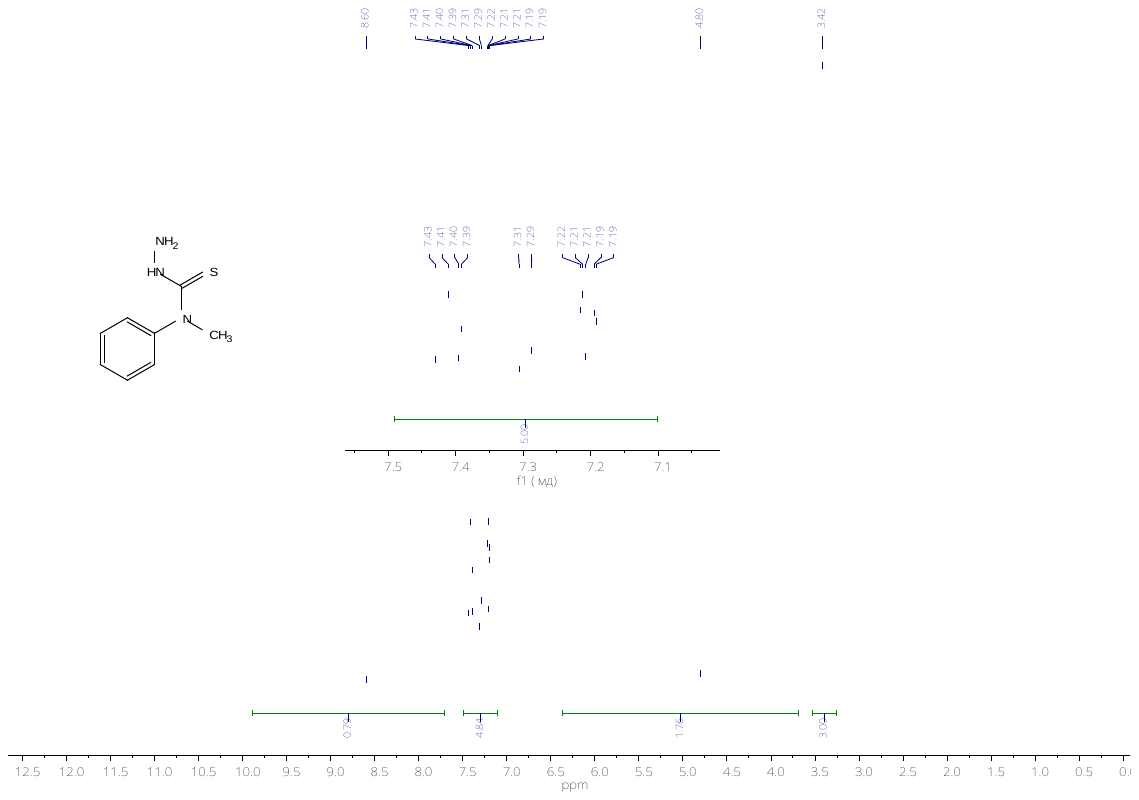

Figure S21. NMR ^1^H spectrum of 4-methyl-4-phenyl-3-thiosemicarbazide (8)

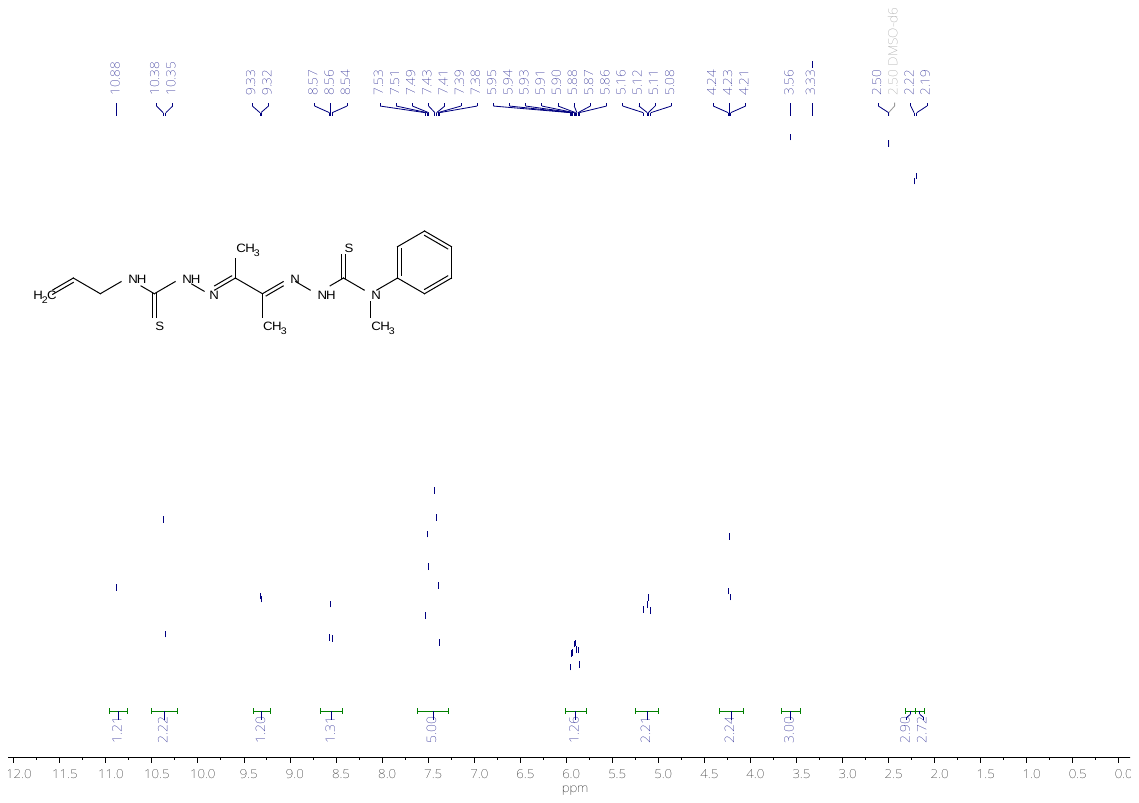

Figure S22. NMR ^1^H (E)-2-((E)-3-(2-(allylcarbamothioyl)hydrazono)butan-2-ylidene)-N-methyl-N-phenylhydrazinecarbothioamide (9a)

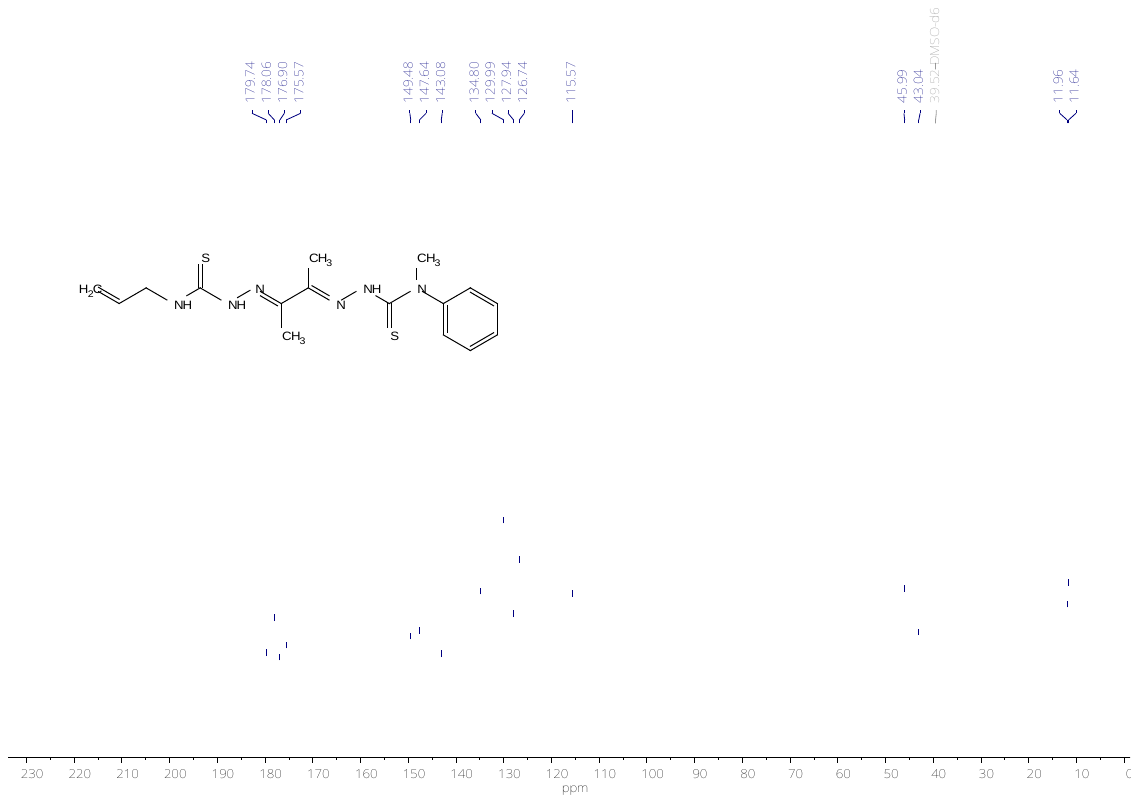

Figure S23. NMR ^13^C spectrum of (E)-2-((E)-3-(2-(allylcarbamothioyl)hydrazono)butan-2-ylidene)-N-methyl-N-phenylhydrazinecarbothioamide (9a)

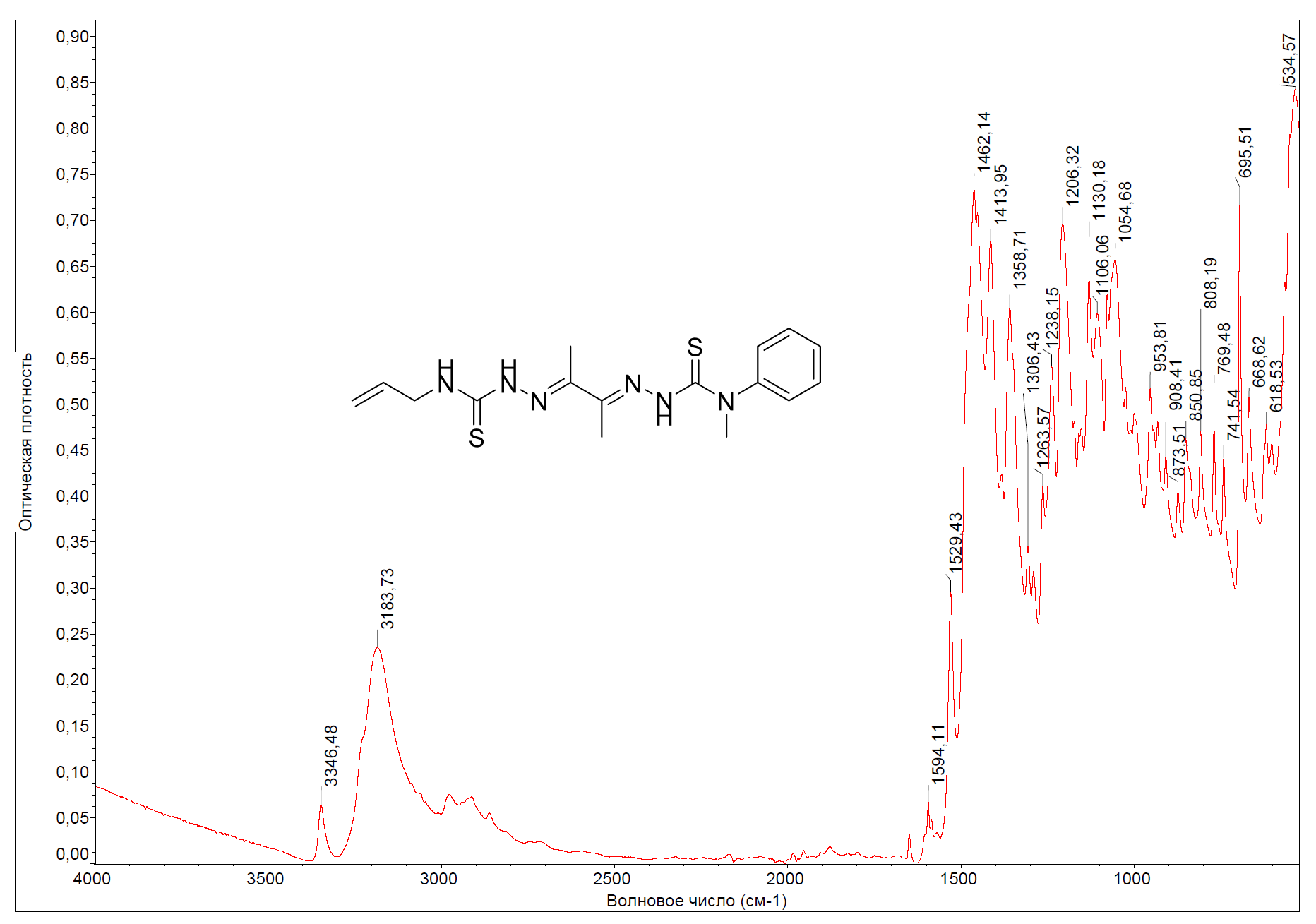

Figure S24. FTIR (E)-2-((E)-3-(2-(allylcarbamothioyl)hydrazono) butan-2-ylidene)-N-methyl-N-phenylhydrazinecarbothioamide (9a)

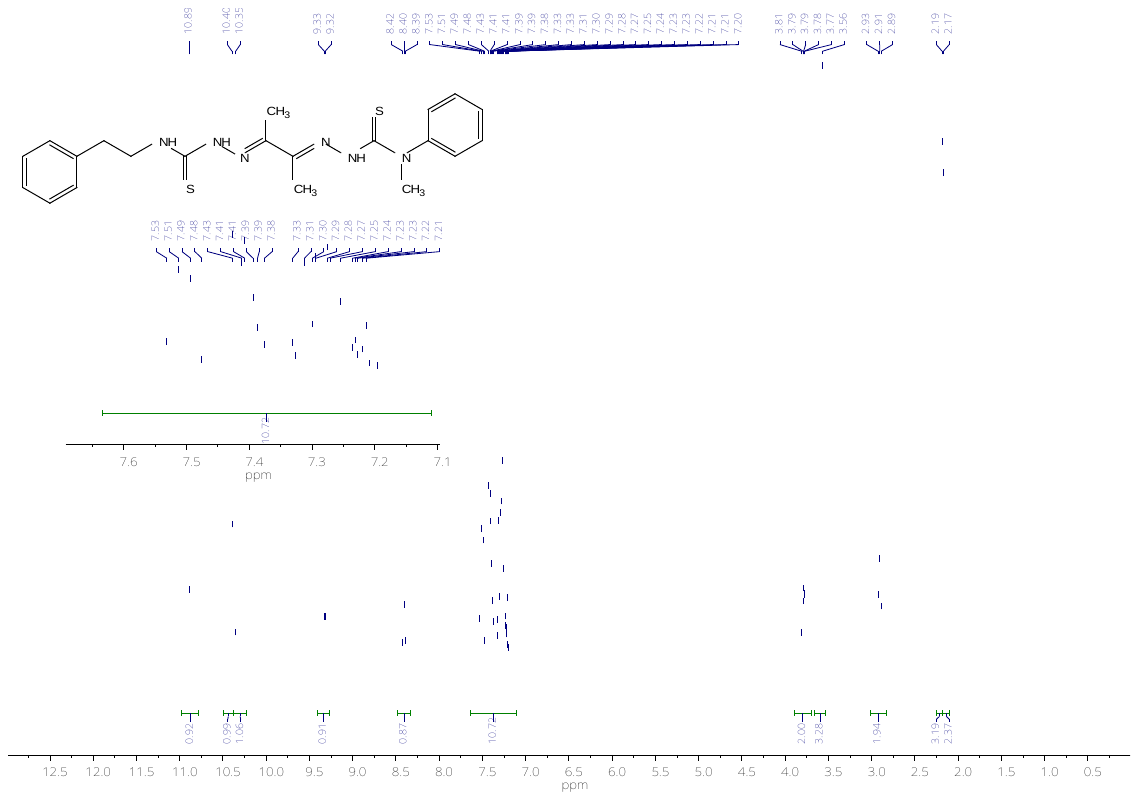

Figure S25. NMR ^1^H spectrum of (E)-2-((E)-3-(2-(phenylethylcarbamothioyl)hydrazono)butan-2-ylidene)-N-methyl-N-phenylhydrazinecarbothioamide (9b)

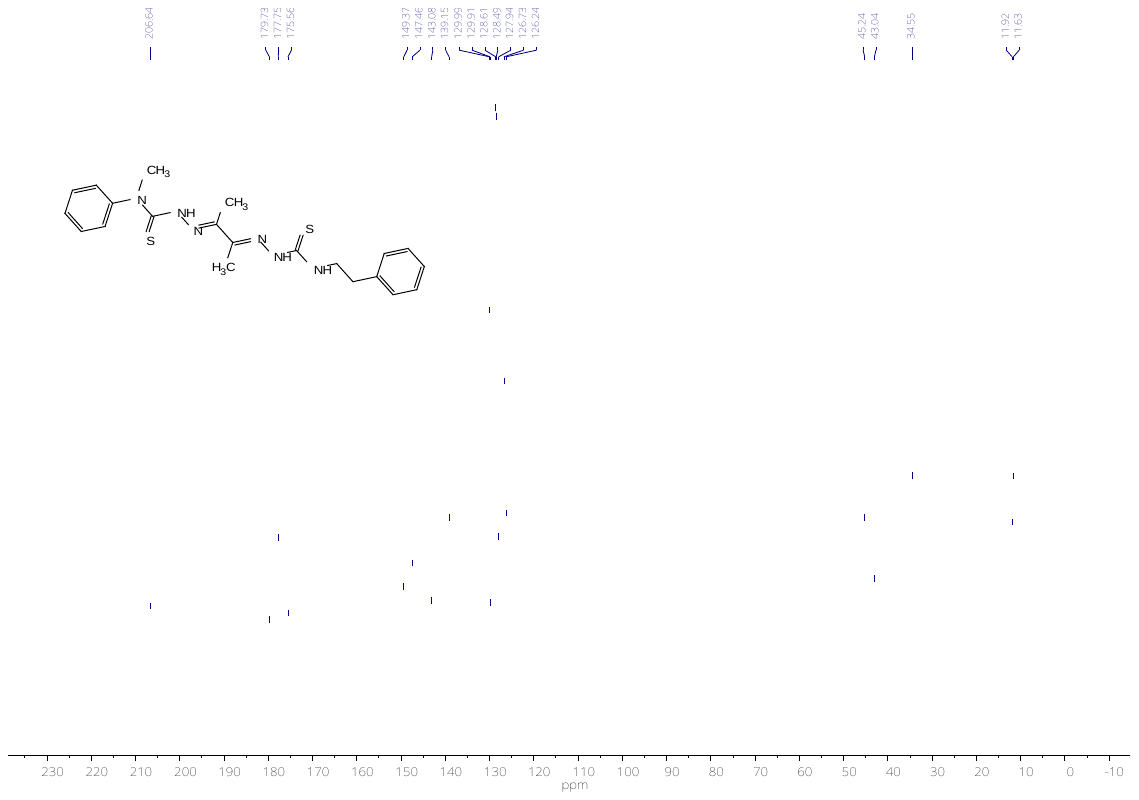

Figure S26. NMR ^13^C spectrum of (E)-2-((E)-3-(2-(phenylethylcarbamothioyl)hydrazono)butan-2-ylidene)-N-methyl-N-phenylhydrazinecarbothioamide (9b)

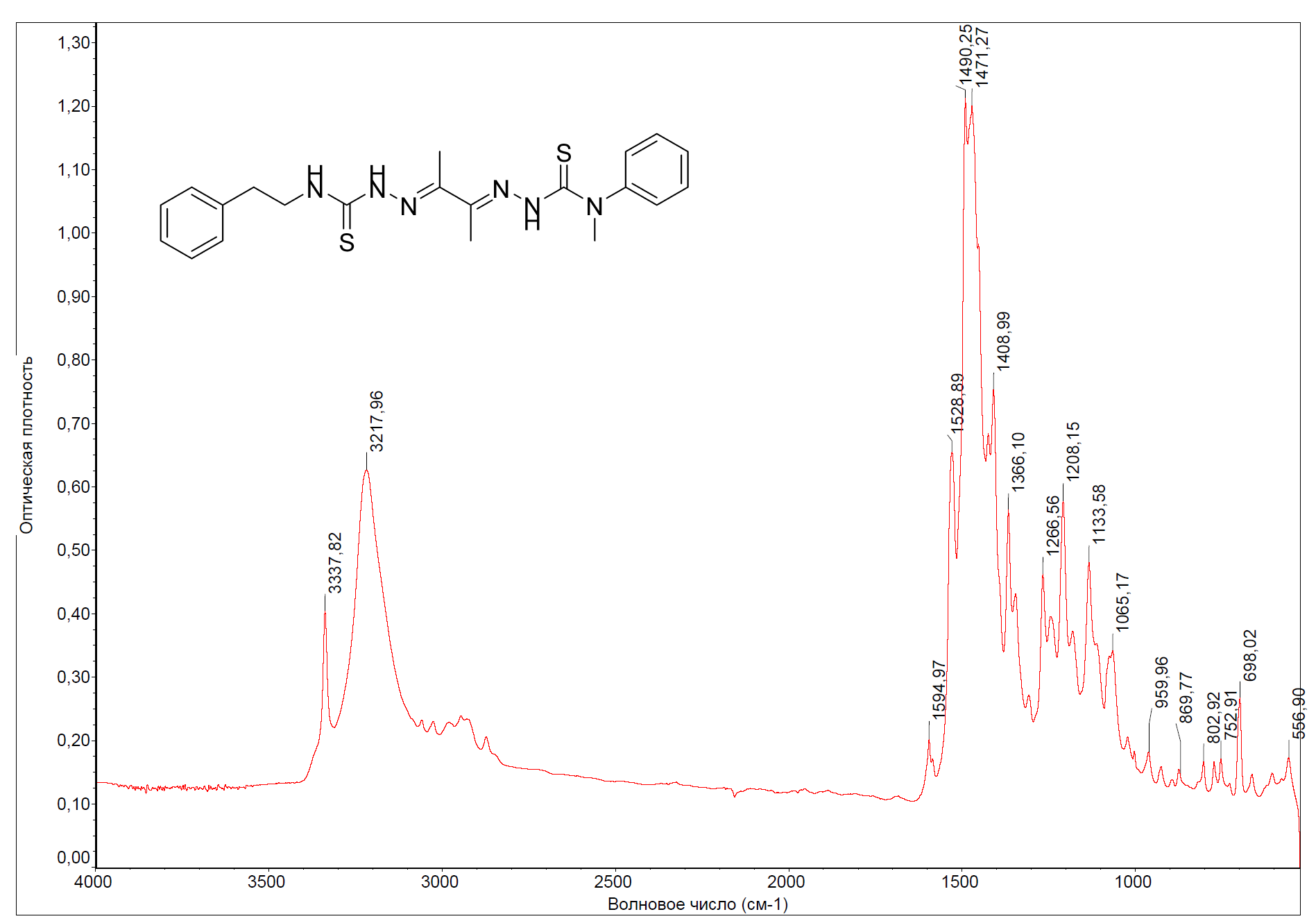

Figure S27. FTIR spectrum of (E)-2-((E)-3-(2-(phenylethylcarbamothioyl)hydrazono) butan-2-ylidene)-N-methyl-N-phenylhydrazinecarbothioamide (9b)

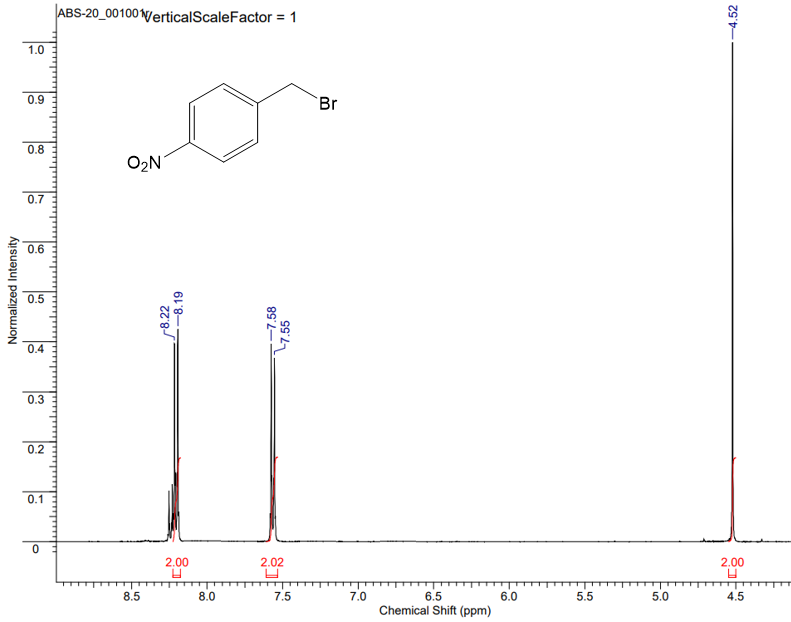

Figure S28. NMR ^1^H spectrum of 4-bromomethylnitrobenzene (10)

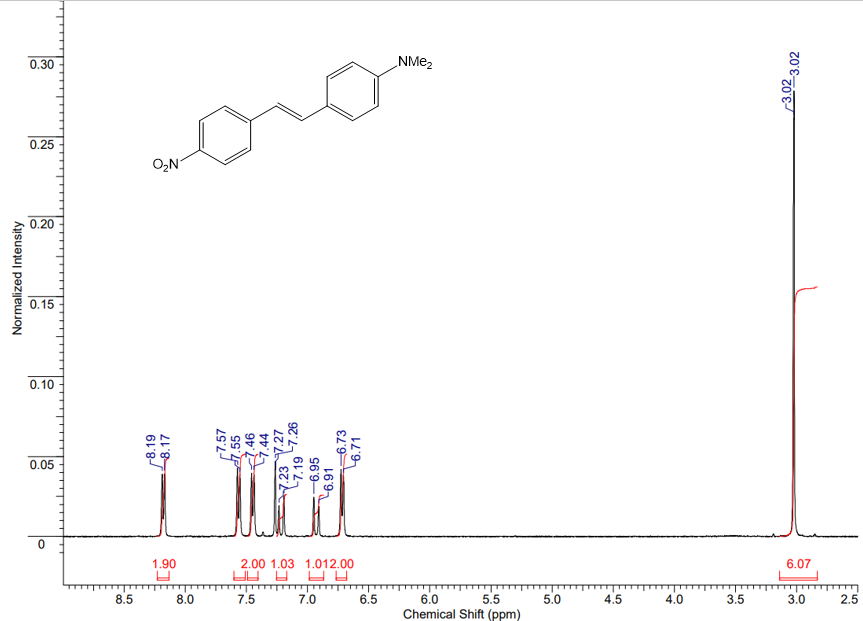

Figure S29. NMR ^1^H spectrum of (E)-4-nitro-4'-dimethylaminostilbene (11)

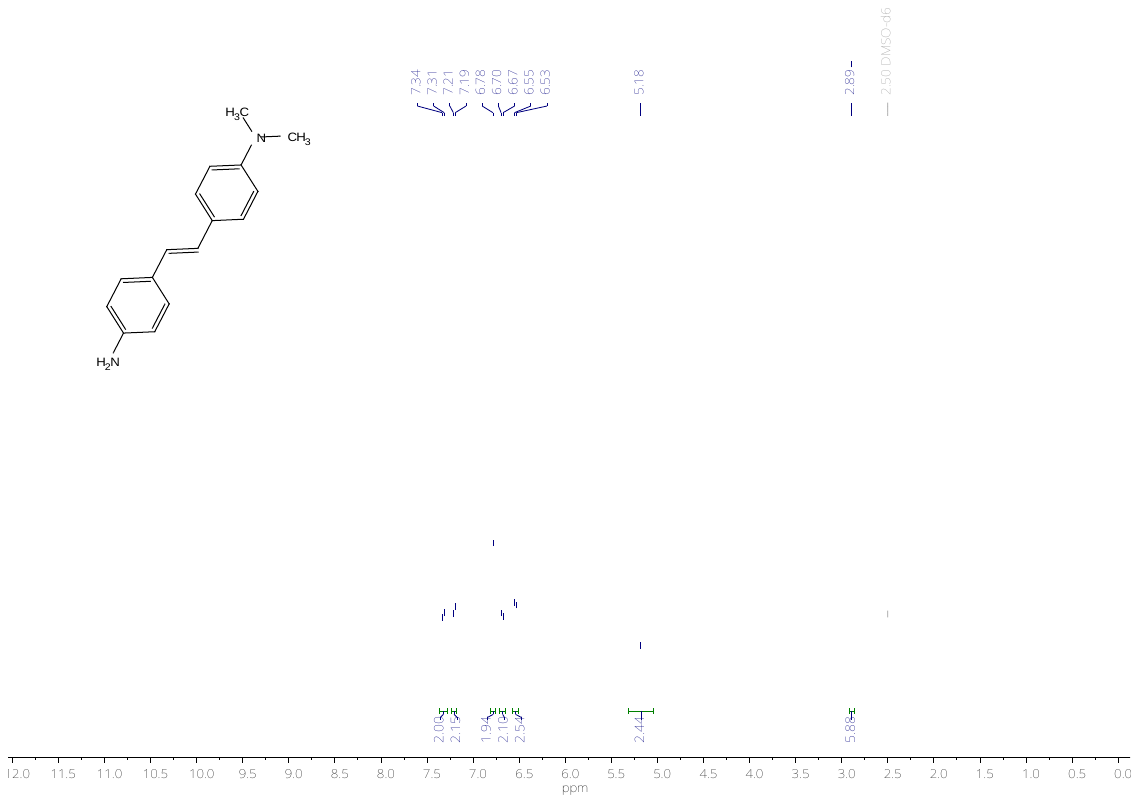

Fig. S30. NMR ^1^H spectrum of (E)-4-amino-4'-dimethylaminostilbene (12)

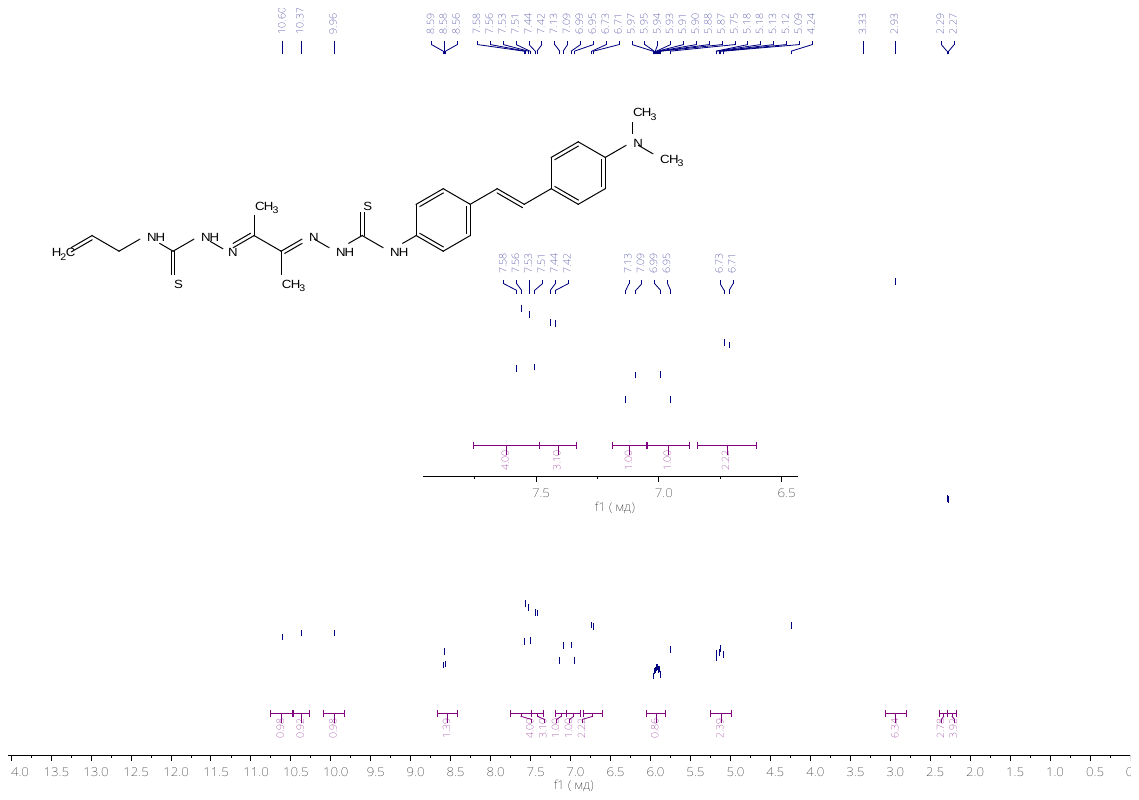

Figure S31. NMR ^1^H spectrum of (E)-N-allyl-2-((E)-3-(2-((4-((E)-4-(dimethylamino)styryl)phenyl) carbamothioyl)hydrazono)butan-2-ylidene)hydrazinecarbothioamide (13a)

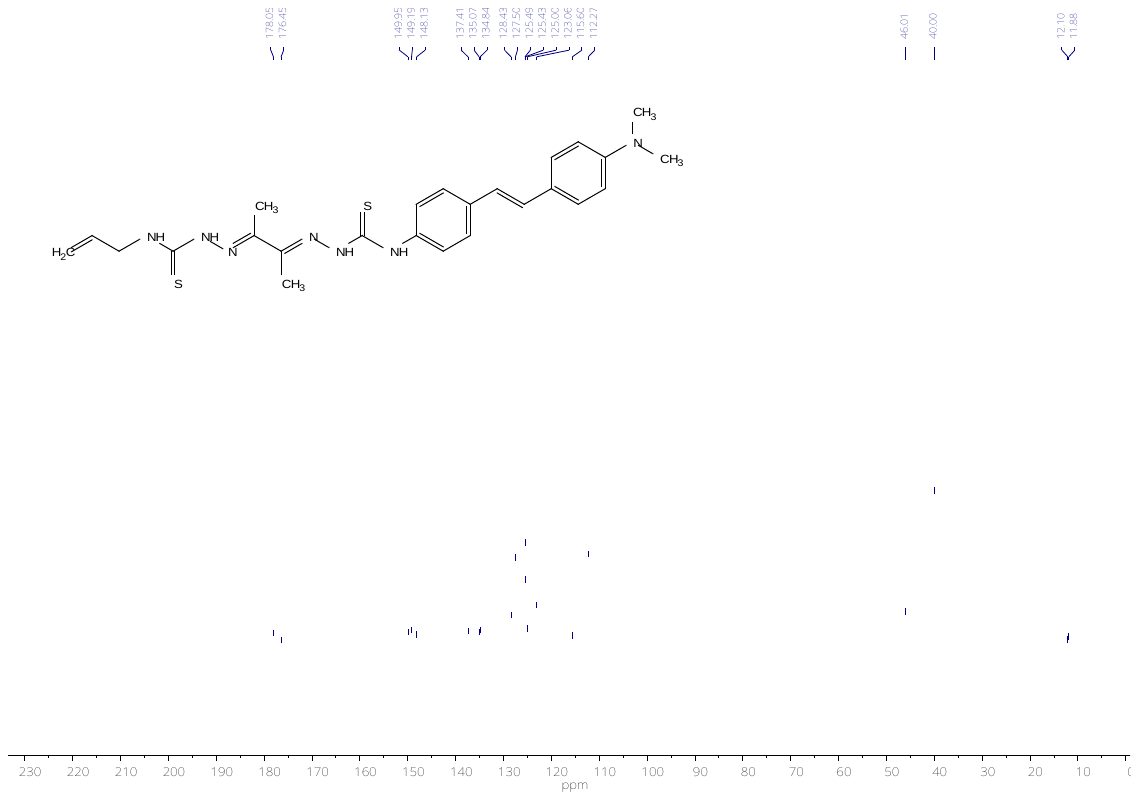

Figure S32. NMR ^13^C (E)-N-allyl-2-((E)-3-(2-((4-((E)-4-(dimethylamino)styryl)phenyl) carbamothioyl)hydrazono)butan-2-ylidene)hydrazinecarbothioamide (13a)

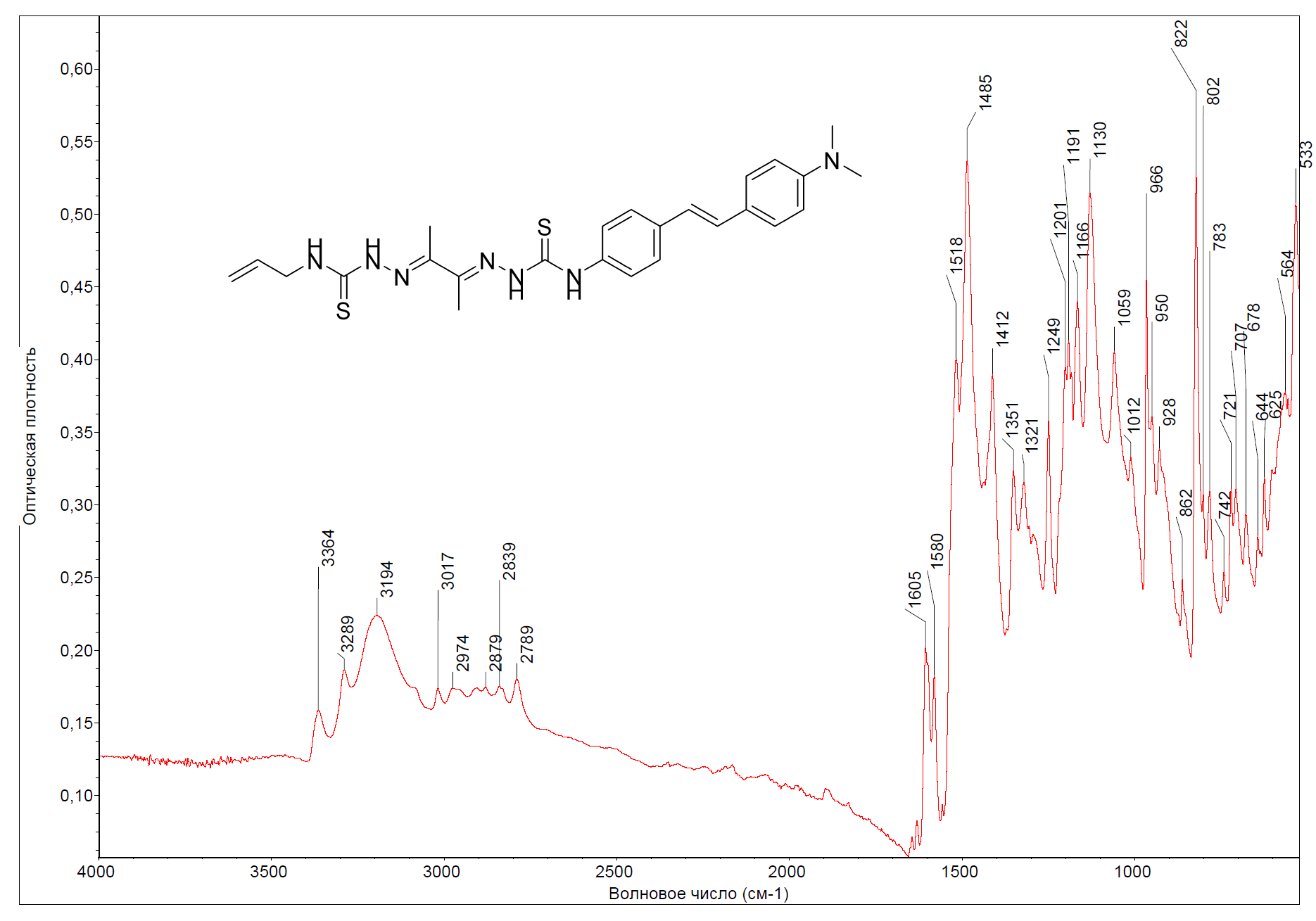

Figure S33. FTIR (E)-N-allyl-2-((E)-3-(2-((4-((E)-4-(dimethylamino)styryl )phenyl)carbamothioyl) hydrazono) butan-2-ylidene)hydrazinecarbothioamide (13a)

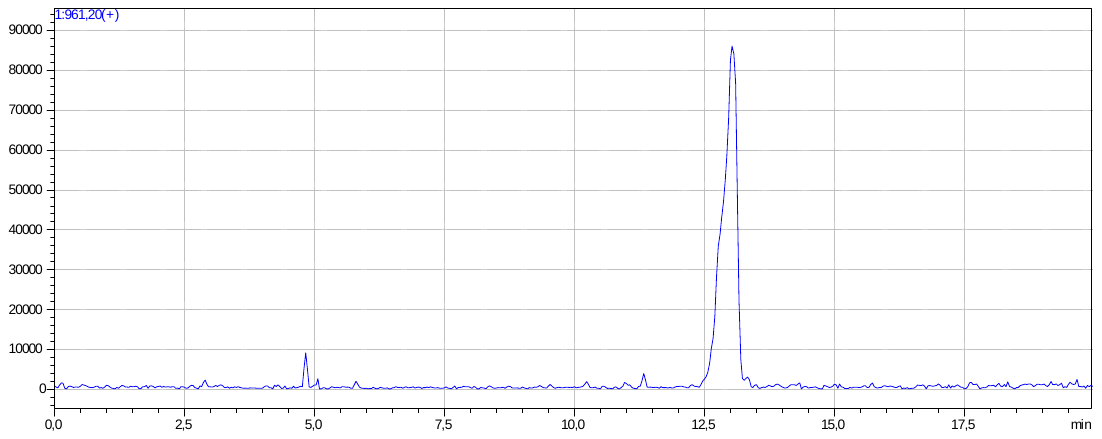

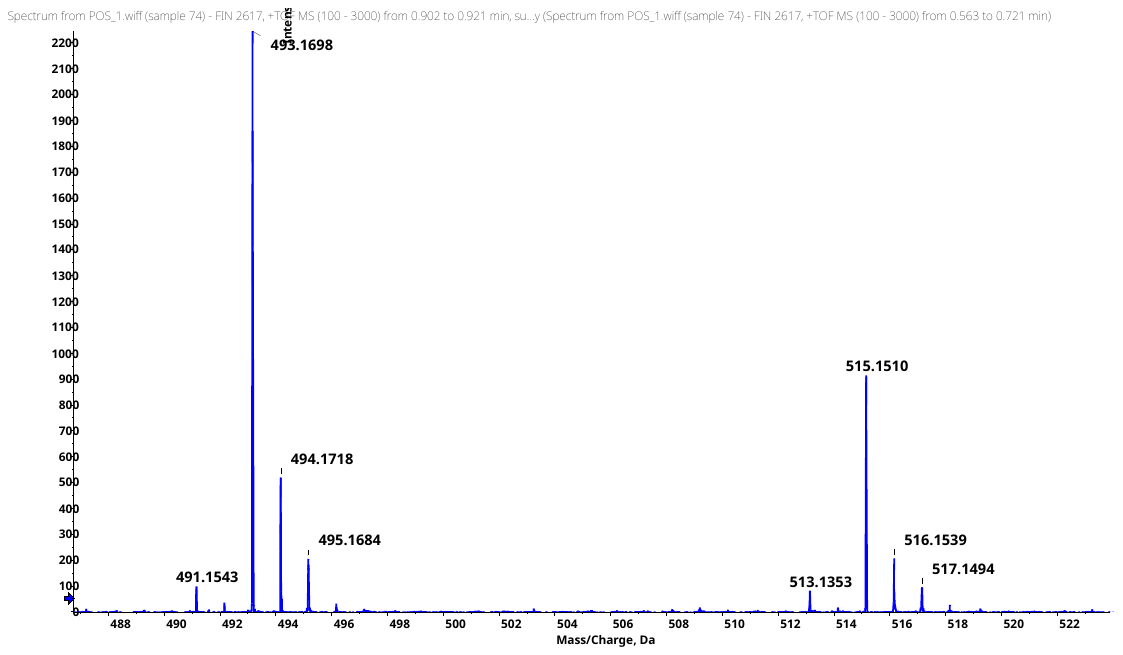

Figure S34. HR-MS data on (E)-N-allyl-2-((E)-3-(2-((4-((E)-4-(dimethylamino)styryl)phenyl) carbamothioyl)hydrazono)butan-2-ylidene)hydrazinecarbothioamide (13a)

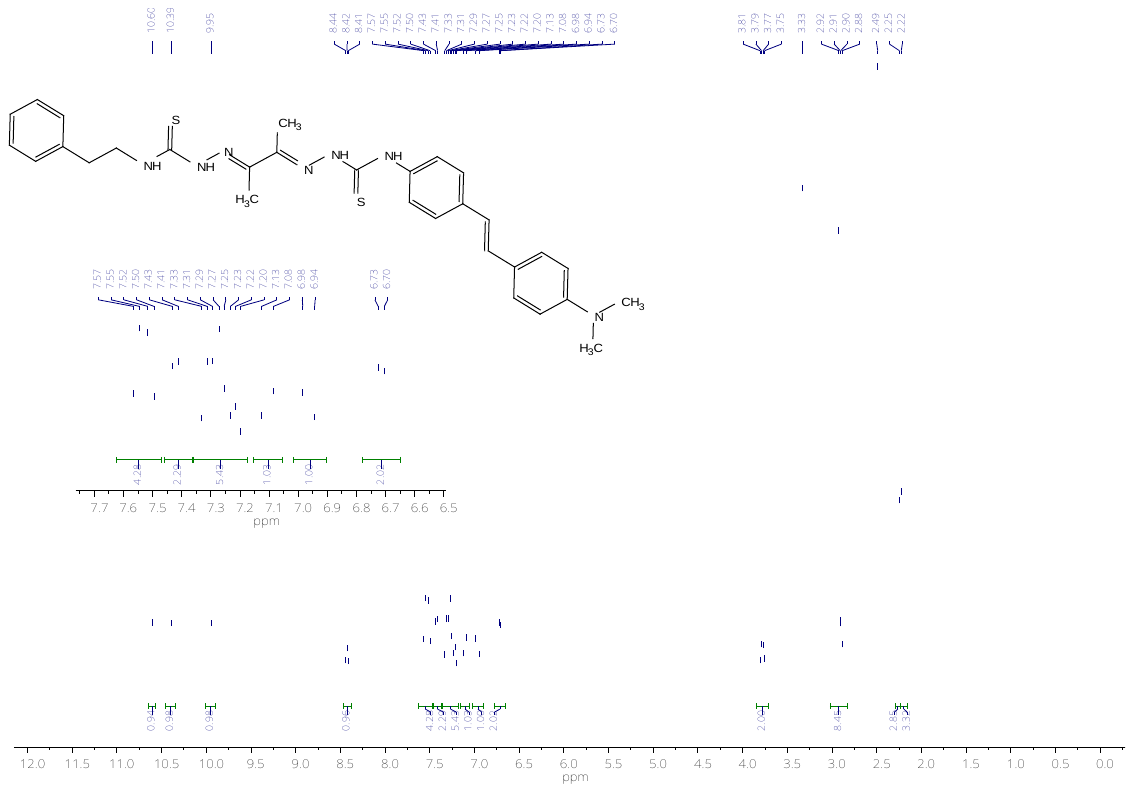

Figure S35. NMR ^1^H spectrum of (E)-N-phenylethyl-2-((E)-3-(2-((4-((E)-4-(dimethylamino) styryl)phenyl) carbamothioyl)hydrazono)butan-2-ylidene)hydrazinecarbothioamide (13b)

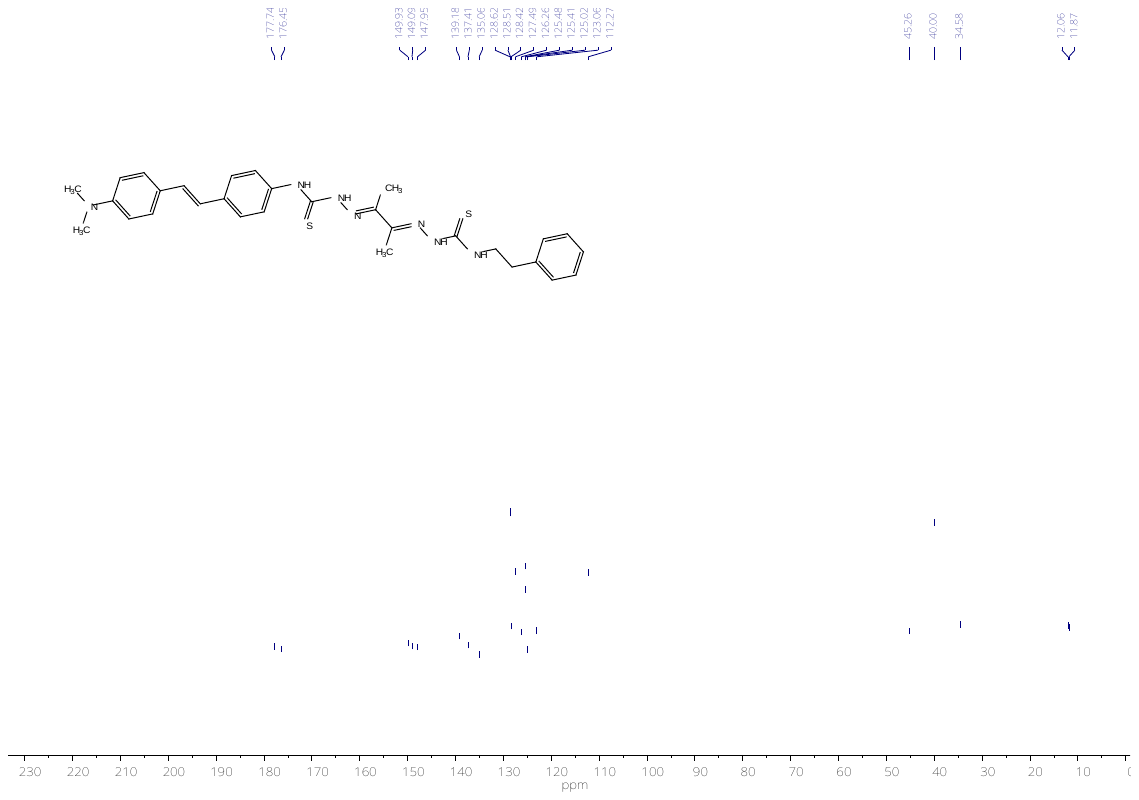

Figure S36. NMR ^13^C spectrum of (E)-N-phenylethyl-2-((E)-3-(2-((4-((E)-4-(dimethylamino) styryl)phenyl) carbamothioyl)hydrazono)butan-2-ylidene)hydrazinecarbothioamide (13b)

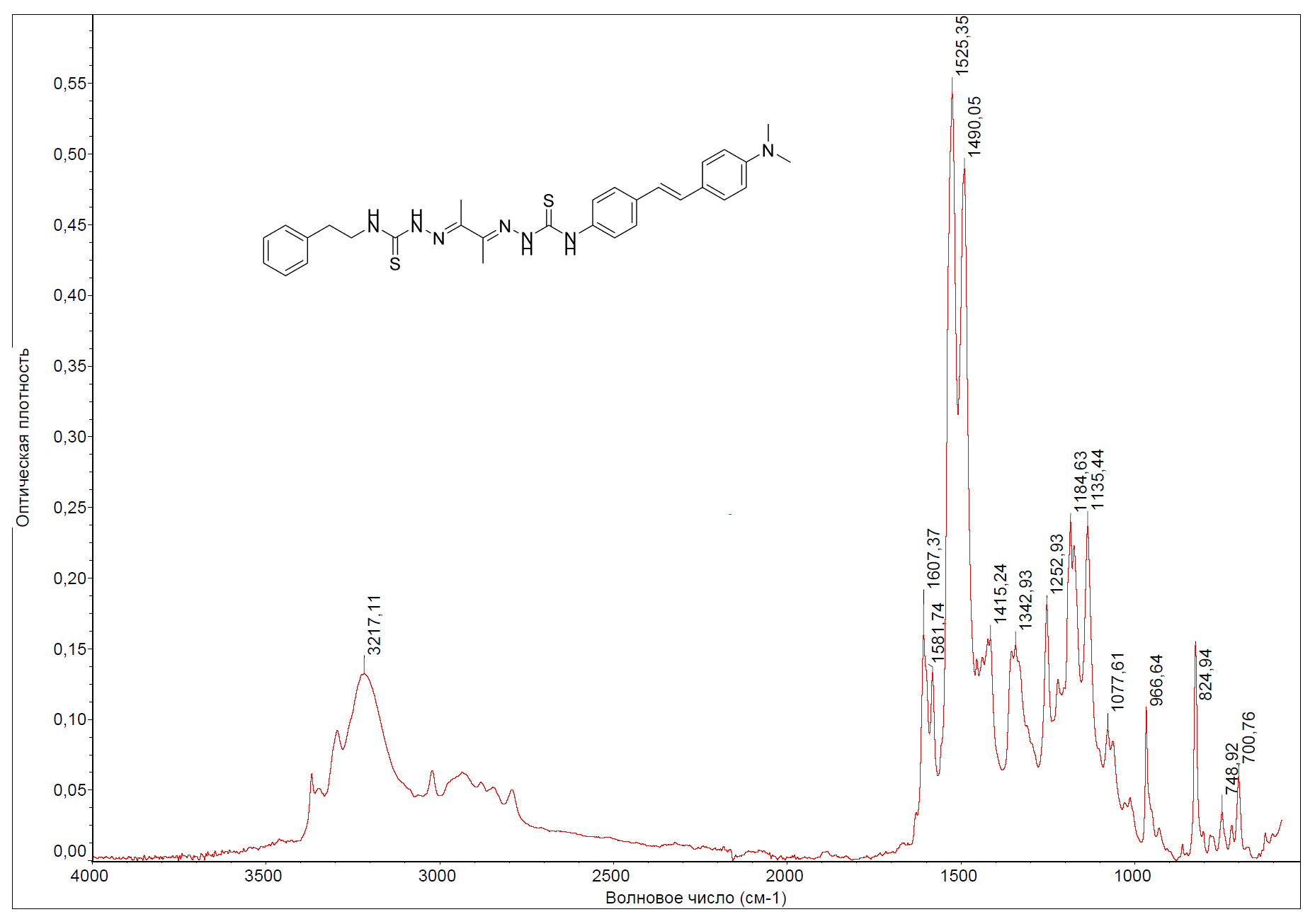

Figure S37. FTIR spectrum of (E)-N-phenylethyl-2-((E)-3-(2-((4-((E)-4-(dimethylamino) styryl)phenyl) carbamothioyl)hydrazono)butan-2-ylidene)hydrazinecarbothioamide (13b)

### Amyloid preparation and cytotoxicity measurements

**Amyloid preparation**

A preparation of synthetic peptide of 40 amino acids β-amyloid Aβ_40_ (PepMic, China) was carried out using standard technology [1]. For this, 1 mg of powder was dissolved in 1,1,1,3,3,3-hexafluoro-2-propanol (Sigma, USA) on ice to a final peptide concentration of 1 mM in a glass vial; after dissolution, the solution was incubated 1 hour at room temperature to obtain peptide monomers. Then the vial with the peptide was placed back on ice for 5-10 min and the resulting solution was aliquoted into microtubes. Next, the tubes were left open for evaporation of 1,1,1,3,3,3-hexafluoro-2-propanol, the remaining alcohol was evaporated on a rotary evaporator for 1 hour, and the resulting films were stored at -70 °C.

**Cytotoxicity assay**

For the study IC_50_ of substances and Cu-ions effect, cytotoxicity assay was performed using HepG2 and SH-SY5Y cell lines (ATCC) and the MTS reagent (Promega). The standard research protocol consisted of the following steps: 10,000 cells were planted in the wells of a 96-well plate in DMEM/F12-based growth medium with 10% FBS, 2 mM L-glutamine, 100 units/ml penicillium, 100 units/ml streptomycin. After 24 hours of incubation at 37°C, 5% CO_2_, the growth medium was replaced with a medium containing the test substances from 0.4 to 100 uM concentrations (DMSO content 0.1 5 and less) with and without CuCl_2_ in the same concentrations and incubated for 48 hours.

For studying substances in Aβ_1-40_ presence, 20,000 SH-SY5Y cells were planted in the wells of a 96-well plate in DMEM/F12-based growth medium with 10% FBS, 2 mM L-glutamine, 100 units/ml penicillium, 100 units/ml streptomycin. After 24 hours of incubation at 37°C, 5% CO_2_, the growth medium was replaced with mix, which consisted of a medium without FBS, the test substances with and without CuCl_2_ in the same concentration and Aβ_1-40_ or fibrils at 10 uM concentration and incubated for 48 hours. Fibrils with and without CuCl_2_ were performed from Aβ_1-40_ in PBS for 24 h at 37^o^C following the described protocol [2].

Afterwards, the medium with the drugs was replaced with the medium with the MTS reagent, incubated for 4 hours, and the optical density values in the wells were measured at 490 nm on a microplate spectrophotometer (Thermo Scientific Multiskan GO).

### Cytotoxicity data

1. Cytotoxicity on HepG2 cells

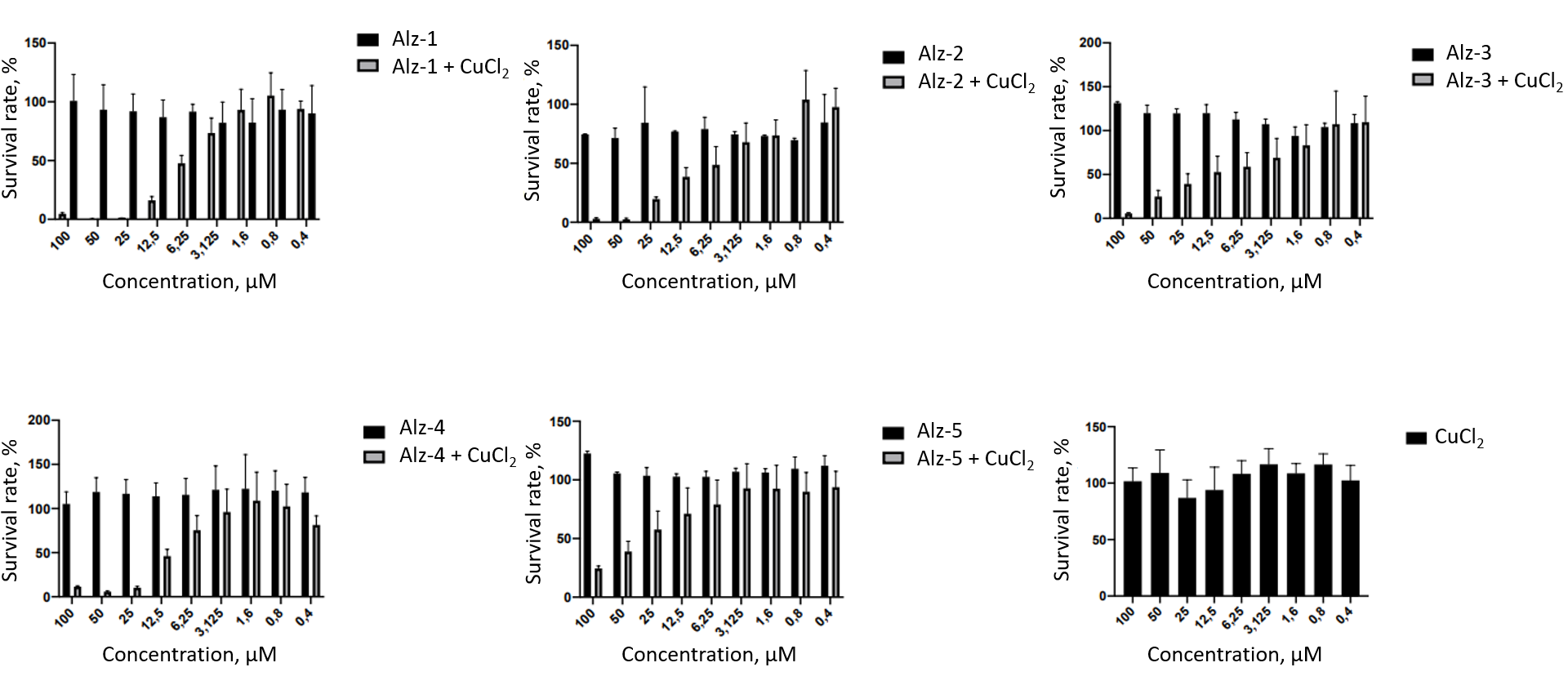

Figure S38. Toxicity data on HepG2 cells.

2. Cytotoxicity on SH-SY-5Y cells

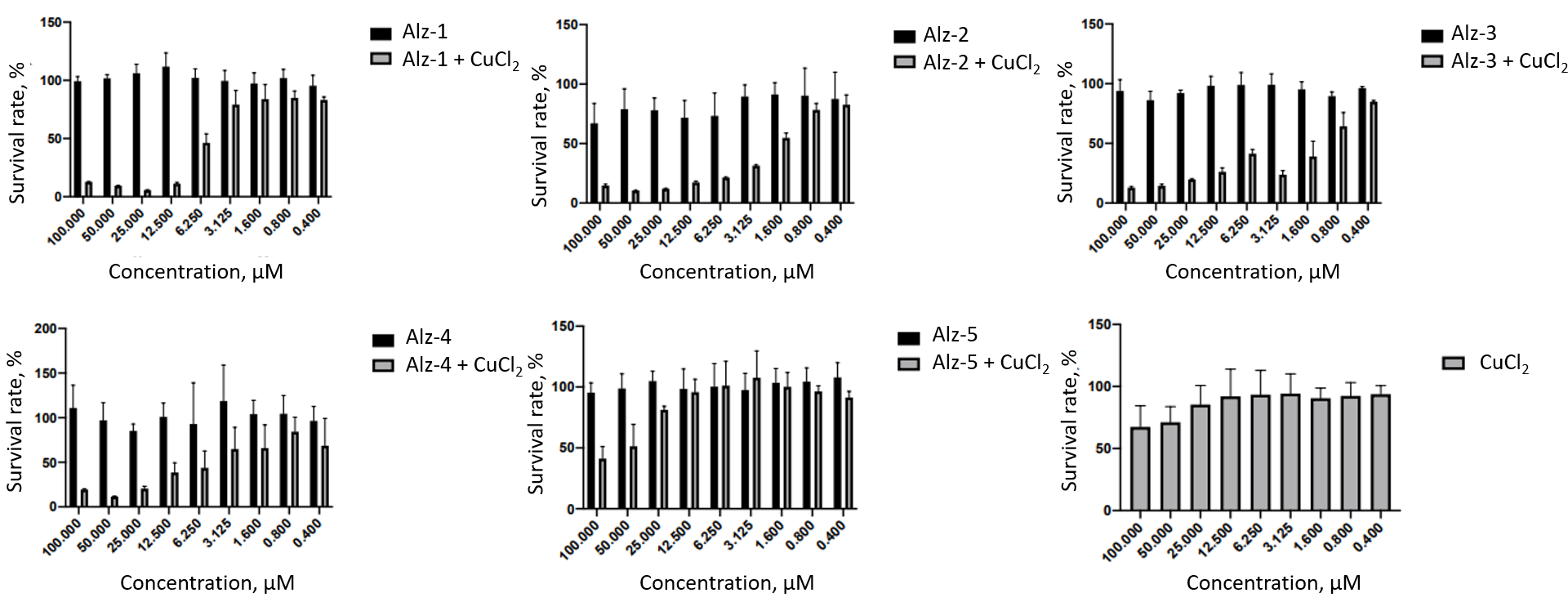

Figure S39. Toxicity data on SH-SY-5Y cells.

### Amperometric ROS measurements

**Method description**

The total ROS concentration was determined by the amperometric method using Pt-nanoelectrodes. Commercially available disk-shaped carbon nanoelectrodes isolated in quartz (ICAPPIC Limited, UK) with diameters 60–100 nm were used to prepare Pt nanoelectrodes. Firstly, the carbon surface was etched in a 0.1 M NaOH, 10 mM KCl solution during 40 cycles of 10 seconds (from 0 to +2200 mV) to create nanocavities. Further electrochemical deposition of platinum in nanocavities was achieved by cycling from 0 to 800 mV with a scan rate of 200 mV/s for 4 to 5 cycles in 2 mM H_2_PtCl_6_ solution in 0.1 M hydrochloric acid. Cyclic voltammetry from -800 to 800 mV with a scan rate of 400 mV/s in a 1 mM solution of ferrocene in methanol in PBS was used to control the electrode surface at all stages of fabrication. Prior to the measurements, each platinum nanoelectrode was calibrated using a series of standard H_2_O_2_ solutions at a potential of +800 mV vs Ag/AgCl. Preparation of Pt-nanoelectrodes has been described in detail elsewhere [3,4].

SH-SY5Y (3 × 10^5^) cells were seeded in 35 mm Petri dishes and treated on the next day with Aβ_42_ / Alz-5 or Aβ_42_ with Alz-5 simultaneously. Aβ_42_ and Alz-5 were dissolved in DMSO. The final concentration of Aβ_42_ / Alz-5 in the culture medium were 10 µM (Aβ_42_) / 50 µM (Alz-5) with 4 h of incubation time. Untreated cells were used as a control, which was performed at the beginning and at the end of the experiment. Before experiment, attached cells in Petri dishes were washed three times by using Hanks’ Balanced Salt solution to remove the growth media and traces of Aβ_42_ / Alz-5. A nanoelectrode penetrated the cells and measured the oxidation current of hydrogen peroxide. On average, about 15 cells were measured by 2–3 Pt electrodes in independent Petri dishes.

The setup for amperometric measurements included a PC that was connected to a system consisting of an ADC-DAC converter Axon Digidata 1550B (Axon Instruments, USA) and patch-clamp amplifier MultiClamp 700B (Axon Instruments, USA). The working head of the amplifier was fixed on a PatchStar Micromanipulator (Scientifica, UK), which placed near an inverted optical microscope Nikon Eclipse TI-U (Nikon, Japan). A Pt nanoelectrode was fixed by a special holder on the working head of the amplifier. The potential difference between the Pt nanoelectrode and the reference (Ag/AgCl) electrode was recorded with the pClamp 11 software suite (Molecular Devices, USA) and processed with Origin 2018 software.

### Scanning Ion-Conductance Microscopy (SICM)

**Materials and methods**

SICM by ICAPPIC (ICAPPIC ltd, United Kingdom) is used for topography mapping and estimation of Young's modulus of SH-SY5Y cells. Nanopipettes with typical tip radius 45-50 nm was fabricated from borosilicate glass O.D. 1.0 mm, I.D. 0.5 mm (WPI, United Kingdom) by using laser puller P-2000 (Sutter Instruments, USA). Nanopipette radius was calculated by using by following theoretical model [5]:

$$r=\frac{Io}{\pi Vktg(\alpha)}$$

where the half-cone angle α is 3 degrees, κ is 1.35 S m−1 and V is the applied electrical potential of 200 mV.

Scanning procedure performed in Hank’s solution (Gibco, USA). Each experimental point had 20–30 cells scanned. For estimating Young's modulus of living cells [6], a nanopipette was approached to the surface until the ion current through the tip reduced by 2% from its initial value during scanning. A noncontact topographic image was obtained at an ion current decrease of 0.5% and further two images were obtained at an ion current decrease (or set-point) of 1% and 2% corresponding to membrane deformations produced by intrinsic force at each setpoint. Then, Young’s modulus estimated by using following model:

$$E=PA{(\frac{Ssub}{Scell}-1)}^{-1}$$

where *E* is the estimated Young’s modulus, *P* is the applied pressure, A is a constant depending on the nanopipette geometry, and S_sub_ and S_cell_ are the slopes of the current–distance curve observed between the ion current decreases of 1% and 2% at the non-deformable surface (S_sub_ – substrate) and cell surface (S_cell_), respectively.

### Atomic-Force Microscopy (AFM)

**Sample preparation**

Stock solution of β-amyloid peptide in DMSO (1.25 mM) was dissolved in filtered PBS 1x pH 4.0 to a final concentration of 100 μM. 4 solutions were prepared, after which: 1. the first solution was used as a control sample (Aβ_42_); 2. 100 μM Aβ_42_ + 100 μM CuCl_2_ (Aβ_42_ + CuCl_2_); 3. 100 μM Aβ_42_ + 100 μM Alz-5 (Aβ_42_ + Alz-5); 4. 100 μM Aβ_42_ + 100 μM CuCl_2_ + 100 μM Alz-5 (Aβ_42_ + CuCl_2_ + Alz-5). All solutions were carefully resuspended and incubated at 37 °C for 24 hours. After the incubation time, the solutions were thoroughly mixed by vortex and diluted with ultrapure water 10 times to a final concentration of Aβ_42_ 10 μM. For atomic force microscopy scans 10 μL of each sample was placed on freshly cleaved mica and incubated in a Petri dish for 10 minutes for peptide adsorption onto the mica. After the adsorption time, the mica samples were washed with 1 ml of ultrapure water and dried in an argon flow. AFM scans was carried out on an NTEGRA II microscope (NT-MDT SI) in semi-contact mode with silicon cantilevers (1.74 N/m, 90 kHz), scan rates 0.5–0.8 Hz, oscillation amplitude is 20 nm, Gain 0.5–0.7, scan size 1–5 μm, 512 points.
